# Proteome-Wide Bioinformatic Annotation and Functional Validation of the Monotopic Phosphoglycosyl Transferase Superfamily

**DOI:** 10.1101/2024.07.10.602977

**Authors:** Theo Durand, Greg J. Dodge, Roxanne P. Siuda, Hugh R. Higinbotham, Christine A. Arbour, Soumi Ghosh, Karen N. Allen, Barbara Imperiali

## Abstract

Phosphoglycosyl transferases (PGTs) are membrane proteins that initiate glycoconjugate biosynthesis by transferring a phospho-sugar moiety from a soluble nucleoside diphosphate sugar to a membrane-embedded polyprenol phosphate acceptor. The centrality of PGTs in complex glycan assembly and the current lack of functional information make these enzymes high-value targets for biochemical investigation. In particular, the small monotopic PGT family is exclusively bacterial and represents the minimal functional unit of the monotopic PGT superfamily. Here, we combine a sequence similarity network (SSN) analysis with a generalizable, luminescence-based activity assay to probe the substrate specificity of this family of monoPGTs in a bacterial cell-membrane fraction. This strategy allows us to identify specificity on a far more significant scale than previously achievable and correlate preferred substrate specificities with predicted structural differences within the conserved monoPGT fold. Finally, we present the proof-of-concept for a small-scale inhibitor screen (eight nucleoside analogs) with four monoPGTs of diverse substrate specificity, thus building a foundation for future inhibitor discovery initiatives.

**Significance:** Uncovering the function and specificity of enzymes responsible for glycoconjugate biosynthesis traditionally requires a multi-faceted and individually curated approach. This is especially true for bacterial glycoconjugates due to greater monosaccharide diversity and a paucity of established structural information. Here we leverage bioinformatic and in-vitro tools to predict and validate substrate specificity for a unique, exclusively bacterial family of enzymes responsible for the first step in many of these glycan assembly pathways. We further show that this platform is suitable for enhanced functional annotation and inhibitor testing, paving the way for the development of urgently needed antibiotics.

## Introduction

Complex glycoconjugates provide structural and functional borders amongst cells and between cells and their environment. In bacteria, glycans contribute to many aspects of functional and diversity (1, 2) and are thus involved in a range of processes including host-cell evasion, colonization, symbiosis, toxin and biofilm generation. Thus, an understanding of the biochemical machinery that enables the assembly of these cell-surface and secreted glycans is timely. In most glycoconjugate biosynthesis pathways, the initiating step occurs on the cytosolic face of the membrane and involves the transfer of a phospho-sugar from a soluble nucleoside diphosphosugar (NDP-sugar) donor to a membrane-embedded polyprenol phosphate (PrenP) (3). In bacteria, the PrenP is most commonly undecaprenol phosphate (UndP) or decaprenol phosphate (DecP) (4-6). The initial transfer step, catalyzed by a phosphoglycosyl transferase (PGT), generates a Pren-PP-linked monosaccharide which is then elaborated by a series of glycosyl transferases (GTs). The sugar moieties themselves can be elaborated by enzymes that act on the NDP-sugar before assembly or directly on the Pren-PP-glycan (Fig. 1).

**Figure 1:**
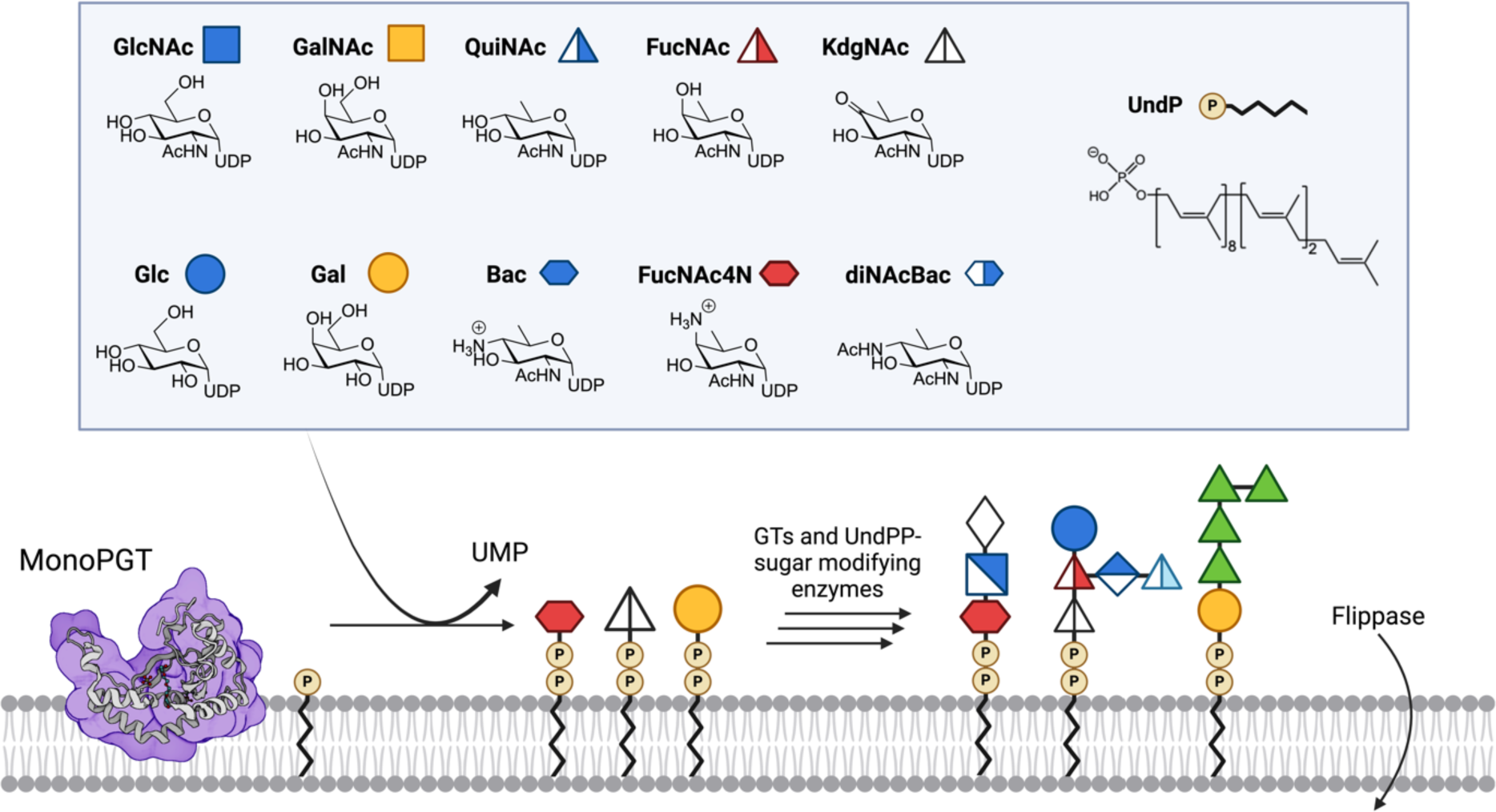
Overview of the monoPGT superfamily activity. Box: Substrates UDP-sugar panel with CFG symbols for Glucose (Glc), Galactose (Gal), *N*-acetylglucosamine (GlcNAc), *N*-acetylgalactosamine (GalNAc), *N*-acetyl-quinovosamine (QuiNAc), *N*-acetyl-fucosamine (FucNAc), Bacillosamine (Bac), 2-acetamido-4-amino-2,4-dideoxy-D-fucose (FucNAc4N), 2,4-diacetamido-2,4,6-trideoxy-D-glucose (diNAcBac) and 4-keto-6-deoxy-GlcNAc (KdgNAc), and the bacterial undecaprenol phosphate. Cartoon illustration of SmPGT superfamily member *C. concisus* PglC (PDB ID: 8G1N) showing re-entrant helix embedded in the cytosolic leaflet of the membrane. Representation of selected glycan repeats for *Fusobacterium nucleatum* ATCC 23726, *Streptococcus suis* serotype 9, and *Acinetobacter baumannii* K92 O-antigens (left to right).

There are two PGT superfamilies, the polytopic (polyPGTs) and the monotopic (monoPGTs), which are differentiated by both membrane topology and catalytic mechanism (7-10). PolyPGTs, exemplified by MraY and GlcNAc-1-P-transferase (GPT), are found in all domains of life, where they are involved in prokaryotic early-stage peptidoglycan and eukaryotic N-linked glycoprotein biosynthesis. These enzymes feature a multi-pass transmembrane topology. PolyPGTs are potently inhibited by several nucleoside natural products including tunicamycin and murayamycin D (11, 12). In contrast, the monoPGT superfamily is exclusively prokaryotic and features a structurally-unique catalytic domain which is embedded in one leaflet of the membrane through an unusual but characteristic re-entrant membrane helix. Currently, this superfamily remains underexplored with respect to both substrate specificity and inhibitor development (Fig. 1).

The monoPGT superfamily is ubiquitous in prokaryotic glycoconjugate pathways and initiates the biosynthesis of glycans found in O-antigen, cell wall polysaccharides, teichoic acids, and glycoproteins, amongst others (13, 14). The superfamily is further classified based on the presence or absence of additional domains outside of the catalytic core (15). MonoPGTs encompassing only the catalytic domain are referred to as small (SmPGTs). MonoPGTs with additional domains can belong to either the large (LgPGT) class, which includes an N-terminal four-transmembrane helix bundle and a nucleotide-binding domain N-terminal to the catalytic domain, or the bifunctional (BiPGT) class, which contain additional catalytic domains such as acetyltransferases or glycosyltransferases fused to the N- or C-terminus of the monoPGT core domain (16). MonoPGTs generally show high specificity for their cognate UDP-sugar substrates, however, the molecular determinants that drive this specificity have remained elusive, even given the information afforded by the recently determined structures of both *Campylobacter concisus* PglC (17), a SmPGT and *Salmonella enterica* WbaP (18), a LgPGT.

Prior work from our laboratory has provided robust PGT assays (19), advanced our knowledge of the catalytic mechanism of monoPGTs (10), provided the first experimentally-determined structures of monoPGTs (17, 18), and identified sequence fingerprints associated with the use of a highly modified UDP-sugar substrate, UDP-*N,N*-diacetylbacillosamine (UDP-diNAcBac) (20). Despite this, both the limited availability of non-standard UDP-sugar substrates and the lack of high-throughput assays represent major hurdles in establishing a more complete understanding of the substrate specificity of the monoPGT superfamily. Further, the membrane-bound nature of the monoPGTs makes these enzymes difficult targets to purify and stabilize after recombinant expression, which has necessitated customized protocols for expression, detergent solubilization, and handling. Analysis of the monoPGT superfamily, either at the bench or bioinformatically, as a whole remains a significant challenge.

Sequence similarity networks (SSNs) provide a graph-based method to analyze large amounts of sequence data, by generating all-to-all alignments where a node (protein sequence) is connected to all other nodes by an edge only if their similarity is above a specified cut-off (21, 22). Thus, SSNs represent clusters of protein sequences that are more ‘similar’ to members of their own cluster than to members of other clusters. Upon optimization of the cut-off value (23), proteins separate into isofunctional clusters, allowing for targeted analysis of functionally-related sequences. Such methods have been shown to be powerful for large-scale sequence analysis (22, 24). An SSN has been previously reported for a representative set of monoPGT sequences from UniProt, which was exhaustively curated through manual approaches (7). This network was very informative but overrepresented by LgPGTs, which complicated the analysis of the resulting clusters due to the presence of the auxiliary domains. Additionally, at a similarity cutoff useful for analyzing the superfamily, the majority of the SmPGTs were segregated into singleton nodes, making it challenging to draw any conclusions as to their relationships.

At the same time, facile methods to confirm predictions of novel substrate specificity were lacking. In the past, two methods were available. The first involved the use of radioactivity-based assays using radiolabeled UDP-sugar substrates that were introduced to PGT cell envelope fractions (CEFs) followed by liquid-liquid phase extraction and scintillation counting to quantify Pren-PP-sugar products (25-27). This time-consuming method is limited by the commercial availability of only the most common radiolabeled NDP-sugars, thus requiring the generation of unique radiolabeled NDP-sugar substrates, but provides high sensitivity. The other established method employs a commercially available luminescence-based assay, UMP-Glo^TM^ (Promega), that enables quantitation of the PGT-transfer reaction product UMP (19). This method has a much higher throughput but requires detergent solubilization of the PGT from isolated CEF, which is often highly detrimental to enzyme activity and stability. The UMP-Glo assay necessitates ultra-pure UDP-sugar substrates owing to potential interference by nucleotide contaminants in the assay sensing mechanism (28).

Herein, we describe a targeted SSN focused on the SmPGT family of the monoPGT superfamily. By mapping PGTs of known substrate specificity onto the resulting clusters, we can computationally predict substrates for uncharacterized PGTs representing the entire superfamily. To biochemically validate these predictions, we develop, benchmark, and apply a modified version of the UMP-Glo assay, which does not require solubilization of target PGTs from the cell membrane, which we term CEF-Glo. Taken together, CEF-Glo and the SmPGT SSN analysis provide a platform for systematically determining substrate specificity of the SmPGT family at much higher throughput and more generally than was previously possible. We apply this toolset to predict and test substrate specificity of several clusters from our SSN. We observe a general preference for the transfer of modified UDP-sugars, providing chemical insight into the logic of glycoconjugate biosynthesis. With new substrate groups assigned, we also leverage recent advances in bioinformatic structural prediction to try and identify conserved structural motifs within the clusters. Finally, we demonstrate that CEF-Glo is a suitable platform for kinetic analysis and inhibitor library screening, widening the reach of our analysis. This approach provides a pipeline to connect sequence, function, and structure at the level of the entire monoPGT superfamily.

## Results

### Sequence similarity network

Network generation was performed using the Enzyme Function Initiative Enzyme Similarity Tool (EFI-EST), with 32,467 SmPGT sequences pooled from the Uniprot database and filtered from the superfamily using a length cutoff to encompass the minimal catalytic domain (21). In order to select the most relevant E-value cutoffs for downstream network analysis and visualization we adapted a recently published methodology for unbiased network analysis (23). Briefly, closeness centrality was calculated across networks with a range of different E-value cutoffs, allowing for a simple and objective graph-based comparison of the quality of the entire set of networks. As the closeness centrality of a network decreases across increasing E-value, the quality of the generated clusters decreases until the network reaches an inflection point and a local maximum. These points of transition represent large-scale rearrangements of the network, which reflect new sequence relationships and improve network quality.

The closeness analysis of the generated SSN revealed E-values for two relevant networks: E-value 61 and 71 (Fig. 2). The E-value 61 network represented the maxima occurring after the most dramatic shift from low to high closeness, whereas the E-value 71 network represented the highest local maximum of the analysis (Fig. S1).

**Figure 2:**
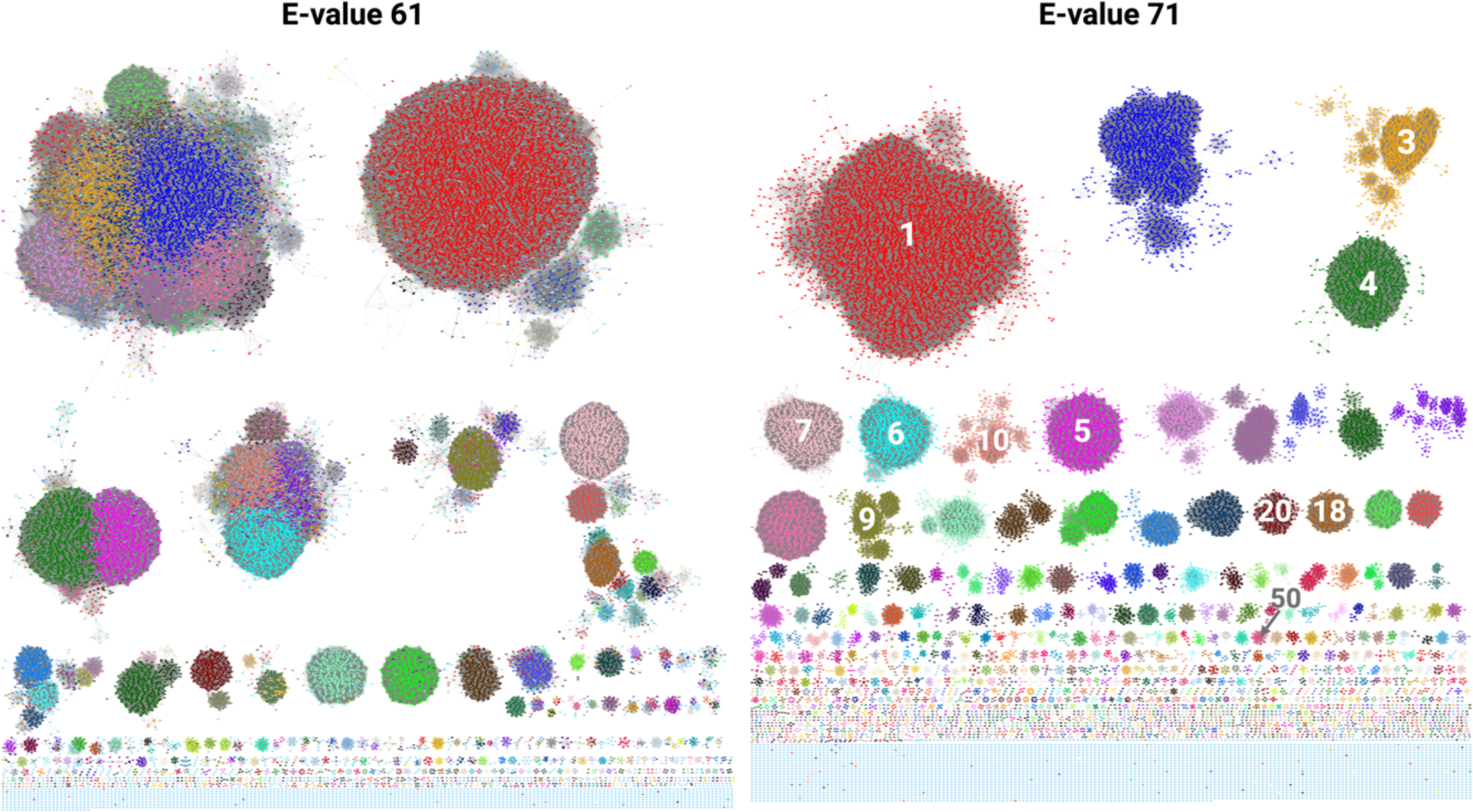
SSNs as selected by Closeness Analysis. Each colored node represents a SmPGT sequence or group of 100% identical SmPGT sequences (100% representative node, repnode). E-61 network colored according to the EFI-EST-applied coloring of network E-71 (descending in order of cluster size, red largest, blue second largest…) to show originating clusters. E-71 The cluster numbers of expressed PGTs (Table 1) are labeled.

**Table 1:** Table of SmPGTs analyzed and expressed in this work. LPS: lipopolysaccharide, CPS: Capsular polysaccharide, CPSA: Capsular polysaccharide A, LTA: Lipoteichoic acid, EPS: Exopolysaccharide. Asterisk indicates *preferred* substrate that is transferred but is not thought to be the physiological substrate.

| Organism | Strain | Gene Name | Glyco-conjugate | Uniprot ID/FASTA | Expression Strain | Predicted UDP-sugar Substrate | E71 SSN |
| --- | --- | --- | --- | --- | --- | --- | --- |
| *Campylobacter concisus* | 13826 | PglC | Protein Glycosylation | A7ZET4 | BL21 pRIL | diNAcBac | 1 |
| *Helicobacter pullorum* | NCTC 12824 | PglC | Protein Glycosylation | E1B268 | BL21 pRIL | diNAcBac* | 1 |
| *Rhizobium etli* | ATCC 51251 CE3 | WreU | O-antigen/LPS | Q2K1T1 | BL21 pRIL | KdgNAc (43) | 5 |
| *Clostridium botulinum* | BL81 | None | Unknown | A0A6B4JHH5 | BL21 pRIL | FucNAc4N | 10 |
| *Streptococcus suis* | Serotype 9 | Cps9F | CPS | Q9RG41 | BL21 pRIL | KdgNAc (63) | 4 |
| *Staphylococcus aureus* | Newman | CAP5M | CPS | P95706 | BL21 pRIL | FucNAc (53) | 50 |
| *Fusobacterium nucleatum* | ATCC 23726 | None | O-antigen /LPS | D5RFH4 | BL21 pRIL | FucNAc4N (64) | 10 |
| *Bacteroides fragilis* | NCTC 9343 | WcfS | CPSA | Q93QV6 | BL21 AI | FucNAc4N (32) | 10 |
| *Streptococcus pneumoniae* | Serotype 2 | None | LTA IV | A0A0H2ZRD5 | BL21 pRIL | FucNAc4N (39, 65) | 6 |
| *Acinetobacter baumannii* | B8300 | ItrA4 | CPS | A0A2I8CW33 | BL21 AI | Gal (66) | 3 |
| *Rhizobium leguminosarum* | bv. trifolii | PssA | EPS | A0A9X5D0X6 | BL21 AI | Glc (67) | 18 |
| *Clostridium difficile* | 630 | CD2783 | PS-II | Q183M0 | C43 | GalNAc (40, 68) | 9 |
| *Vibrio para-haemolyticus* | Serotype O3:K6 | SypR | Structural symbiosis PS | Q87PP3 | BL21 AI | Unknown | 20 |
| *Vibrio vulnificus* | M06-24 | WbfU | CPS | A0A4Q7IER7 | BL21-AI | L-QuiNAc (46) | 7 |

To better understand the relationships between the generated clusters across all E-values sampled for analysis, we manually tracked the largest 50 clusters from the E-71 network across the E-value gradient. By identifying points at which the individual clusters separate, a dendrogram can be built where branching points occur when a cluster separates into two (or more) smaller clusters (Fig. 3). The distance of this branching point to the root represents the E-value at which this separation occurs. By representing the emergence of those clusters across E-values, coupled with information as to substrate specificity, we can identify the points at which isofunctional clusters separate from the aggregate of un-differentiated SmPGT sequences. Further, the E-value at which each cluster separation occurs provides information as to the uniqueness of SmPGT sequence and its connection to function. For example, clusters that separate from the others much earlier in the E-value gradient have distinct sequences that are likely to act on unusual UDP-sugar substrates. On the other hand, clusters that separate from each other later are more likely to perform very similar functions, and therefore act on similar substrates. The dendrogram also allows a broader assignment than would be possible in a single SSN snapshot. It can reasonably be assumed that if two isofunctional clusters separate from a common ancestor at a more stringent E-value, all other clusters separating from that same ancestor will share that same function, with the distinct clusters being the result of separation based on other sequence signatures such as phylogeny, regulatory mechanisms, or interaction partners.

**Figure 3:**
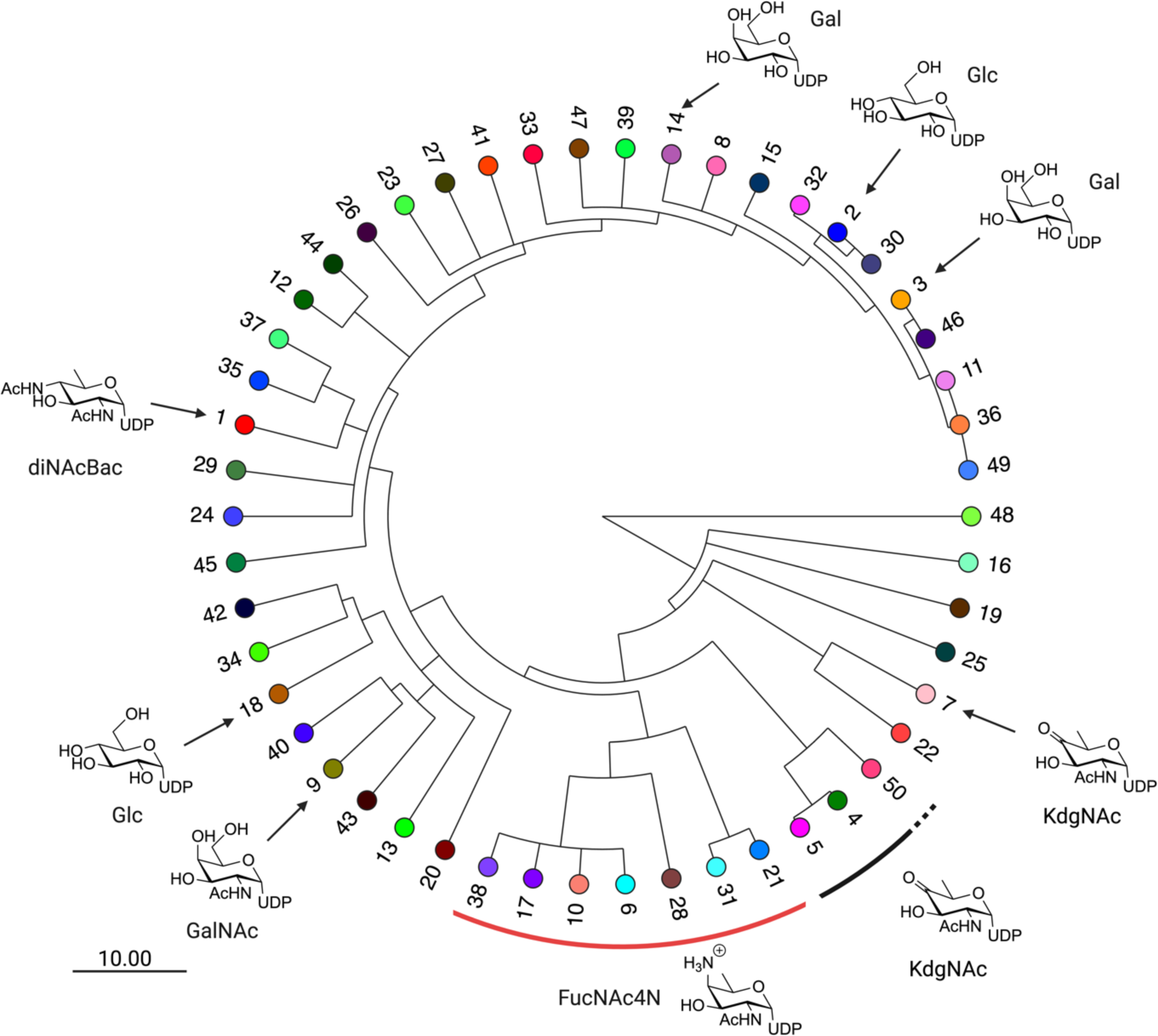
The SSN Dendrogram with leaf nodes that represent clusters in network E-71. The labeling indicates cluster number by descending node count and leaf node coloring corresponds to entries in Fig. 2. Putative and verified substrate assignments are indicated with symbols pointing to the appropriate cluster. Scale bar represents the size of the E-value change encoded in branching point length.

To link the network to a biochemical understanding of the superfamily, we sought to identify putative or experimentally-verified SmPGT – UDP-sugar substrate pairs. Once identified, such pairs would serve as prototypes for identifying the substrates of specific SSN clusters. We identified several characterized SmPGT within the SSN with defined UDP-sugar substrates (Table 1, Table S1). These include PGTs that act on common UDP-sugars, such as UDP-Glc, UDP-Gal, UDP-GlcNAc, and UDP-GalNAc but interestingly, also less common, modified sugars including UDP-QuiNAc, UDP-FucNAc, UDP-FucNAc4N, UDP-diNAcBac and UDP-KdgNAc (Fig. 1, Table S2). By adding those predicted/verified functional assignments to the generated tree we can more easily visualize at what E-value gradient functional separation occurs (Fig. 3). We note that the SSN at E-value 61 forms distinct clusters of sequences transferring highly modified UDP-sugar substrates (UDP-KdgNAc, UDP-FucNAc4N and UDP-diNAcBac) but further E-value optimization is needed separate other isofunctional clusters (Fig. S2). This is reflected in the dendrogram where, from the limited number of clusters with predicted substrate, PGTs with specificity motifs for common UDP-sugars (UDP-Glc and UDP-Gal) are more similar than those for UDP-sugars that have undergone further functionalization and therefore separate later. The points of cluster separation across the SSN may therefore represent the degree for which a unique sequence motif is needed to recognize its cognate substrate.

### Development and Validation of the CEF-Glo assay

Despite the clear separation of SmPGTs into discrete clusters in the SSN, functional validation of putative substrate assignments represents a significant hurdle. Unambiguous determination of substrate specificities requires direct biochemical observation of phosphoglycosyl transfer activity, especially for SmPGTs belonging to clusters with no ‘landmark’ sequence. Standard monoPGT assays like UMP-Glo® and radioactivity-based liquid-liquid extraction methods cannot efficiently scale to assess the number of clusters generated in the SSN. The major bottleneck in scaling the UMP-Glo assay has been the assumed requirement for detergent solubilization and purification of SmPGT targets from the membrane. However, the related ADP-Glo® assay has recently been applied to isolated cell membranes containing target proteins (29). Therefore, we investigated whether we could modify the UMP-Glo assay to reliably function on membrane fractions from *E. coli* overexpressing SmPGTs. This necessitated ensuring a good signal-to-noise ratio across the range of assay conditions (with and without UDP-sugars, with different SmPGTs CEFs) for confident assignment. Further, the purity of the UDP-sugar substrates, particularly those derived through chemoenzymatic methods, would need to be rigorously validated so that there was no interference with the components of the UMP-Glo reagents, which are particularly sensitive to nucleotide impurities.

#### Initial CEF-Glo assay benchmark

PglC, a SmPGT from *Campylobacter concisus*, was selected as the initial test protein for assay development as it met several important criteria: 1) The enzyme is highly overexpressed using a standard expression system (Fig. S3), 2) The purified enzyme is active when assayed using UMP-Glo, requiring low nM enzyme concentrations to produce detectable UMP levels, 3) The substrate scope of this enzyme is well understood, with the native UDP-diNAcBac substrate generated in pure form using established protocols (30).

#### Production of expanded UDP-sugar panel for CEF-Glo assay

With clear success in observing PglC activity in CEFs, we reasoned that a broad set of purified UDP-sugar substrates would enable the rapid paneling of substrate preference for any successfully expressed SmPGT. To generate an expanded set of UDP-sugars for assay, substrates were chosen based on their presence at the reducing end of reported bacterial glycoconjugates (Fig. 1) and the commercial sourcing or availability of characterized synthetic routes. UDP-Glc, UDP-Gal, UDP-GlcNAc, and UDP-GalNAc are commercially available in suitably pure form and require no further purification to be compatible with the assay. For UDP-KdgNAc, UDP-Bac, and UDP-diNAcBac, we have previously published methods for the chemoenzymatic synthesis (30) (Fig. S4). Additionally, with slight modification to the published methodology (31), we utilized sodium borohydride (NaBH_4_) to reduce the carbonyl group of UDP-KdgNAc to afford the two epimeric C4’-hydroxyl reduction products - UDP-FucNAc and UDP-QuiNAc (Fig. S5). UDP-FucNAc4N, a UDP-sugar with an axial C4’ amine group, has a published route for production which relies on modification of UDP-diNAcBac biosynthesis, using the aminotransferase from *Bacteroides fragilis* WcfR (32). The methodology uses a coupled one-pot strategy where UDP-GlcNAc is transformed into UDP-KdgNAc by the *Campylobacter jejuni* redox-dependent dehydratase PglF before being further modified into UDP-FucNAc4N by the action of WcfR. By using previously generated UDP-KdgNAc we modified the protocol to afford the desired product in a single step (Fig. S6). Initial analysis revealed that many of the chemoenzymatically-derived sugars were contaminated with varying amounts of nucleotide impurities, which interfered with the assay readout. Therefore, a general protocol involving treatment of the samples with the promiscuous calf intestinal alkaline phosphatase (CIAP, New England Biolabs), which hydrolyzes phosphate groups from nucleotides while leaving UDP-sugars un-affected was established (33). The final UDP-sugar purifications were carried out by anion-exchange HPLC (Fig. S7).

#### Substrate Screen

With a set of UDP-sugar substrates in hand, we prepared CEFs with overexpressed SmPGTs ranging from well- to poorly-characterized target enzymes (Table 1, Fig. S3). In addition to the level of substrate characterization, the candidate PGTs were chosen based on the SSN and the relevance of the source bacteria with respect to pathogenesis. The generated CEFs were screened against a panel of UDP-sugar substrates (Fig. 4, Fig. S8). Screening results show a high level of specificity for SmPGT substrates and confirm many of the published findings and predictions. The screening data validated the isofunctional character of the identified clusters, predicted new cluster assignments, and highlighted regions where the understanding of SmPGT activity was incomplete. Apart from showing clear activity with the predicted cognate substrate, we were able to reveal interesting patterns of specificity based on chemical structures. For example, SmPGTs from *S. pneumoniae* and *F. nucleatum*, with the cognate substrate UDP-FucNAc4N, also transferred UDP-Bac, a chemically similar substrate with a C4’ primary amine group (Fig. 1 and Fig. 4), albeit to a lesser extent. In contrast, the *C. difficile* SmPGT is highly selective for the cognate substrate, UDP-GalNAc, but less active with UDP-sugar substrates with C4’-axial hydroxyl groups, UDP-Gal and UDP-FucNAc (Fig. 4). The promiscuity toward UDP-FucNAc is particularly interesting given that the sugar is unlikely to be present in the organism and therefore there would be no evolutionary pressure to select against it. These emerging patterns, only now visible by testing with a wide prokaryotic-specific UDP-sugar panel, are intriguing, and may reflect different recognition mechanisms echoed in variation of the residues involved in selecting cognate UDP-sugar substrates.

**Figure 4:**
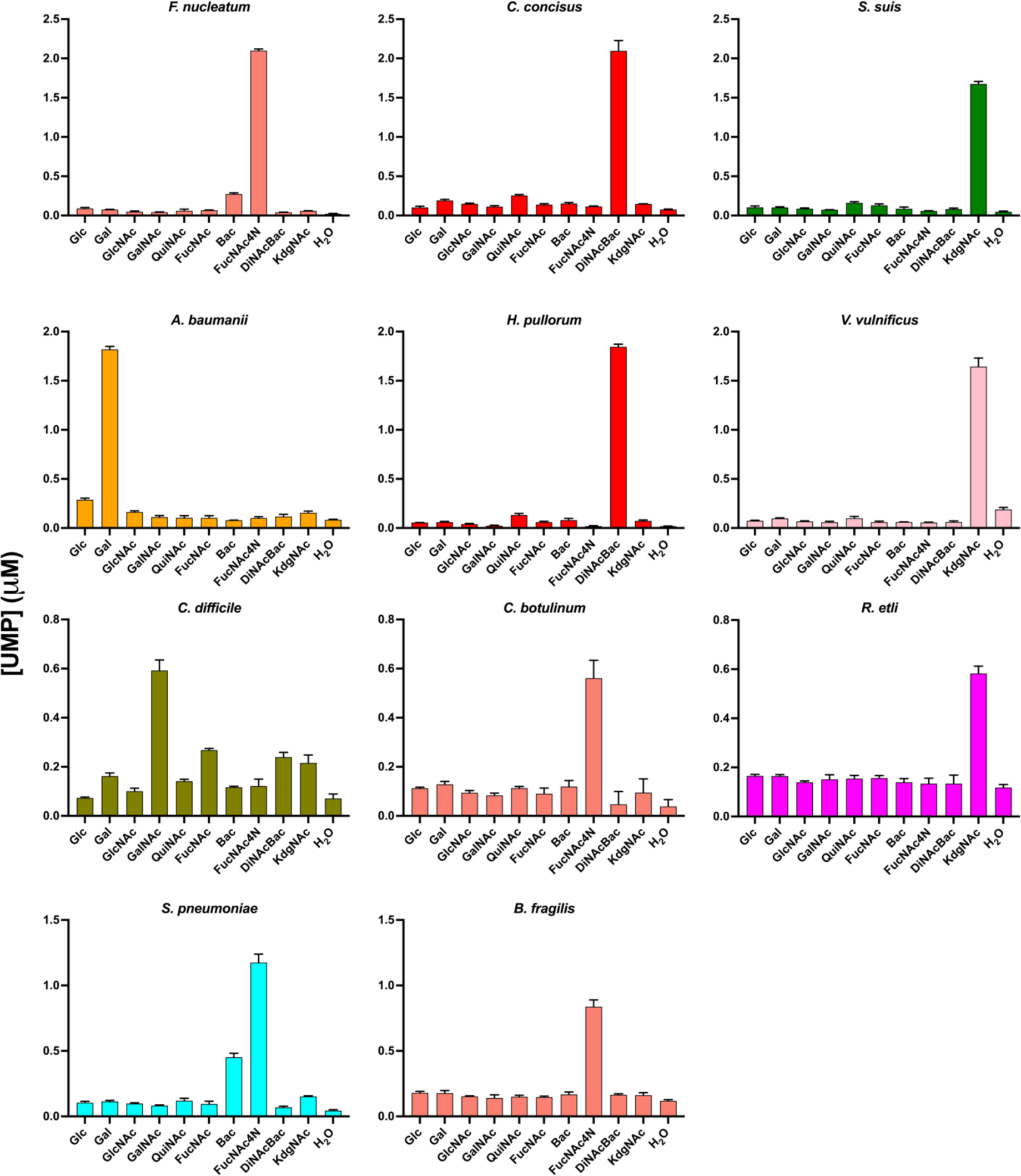
The CEF-Glo substrate screen results with selected monoPGTs. The graph coloring is taken from SSN E-71 cluster coloring. Luminescence measurements in the CEF-Glo assay are shown as the average of triplicate readings converted to [UMP] (µM) using a standard [UMP] curve.

With these annotations, we also leveraged new bioinformatic tools to investigate other features of the clusters. The Foldseek 3Di alphabet encodes AlphaFold-predicted structural information into a sequence that can be aligned and analyzed alongside the amino acid sequence (34, 35). MonoPGTs have a conserved catalytic core and predicted monotopic membrane-associated structure (7), which is reflected in the SSN. However, we were intrigued to see if we could extract further structural features that could be correlated with isofunctional clusters. We used the Foldseek 3Di encoding to compute and compare structural alignments of AlphaFold predicted SmPGT structures across several of the best-characterized clusters (Fig. 5, Fig. S9). By comparing the 3Di alignment from individual clusters to the whole network, we were able to identify potential UDP-sugar specific structural motifs. Notably, two structural features of *C. concisus* PglC that were previously identified as signatures of UDP-diNAcBac specificity, the mobile loop and aromatic box (Fig. S9), are highlighted (20). Analogous regions of other clusters show similarly-conserved structural features that may reflect specificity. Investigating these fine-grained differences in structure at a large scale can provide new insight into structural and chemical aspects of ligand interactions to aid in substrate identification, inhibitor development, as well as experimental structural characterization efforts.

**Figure 5:**
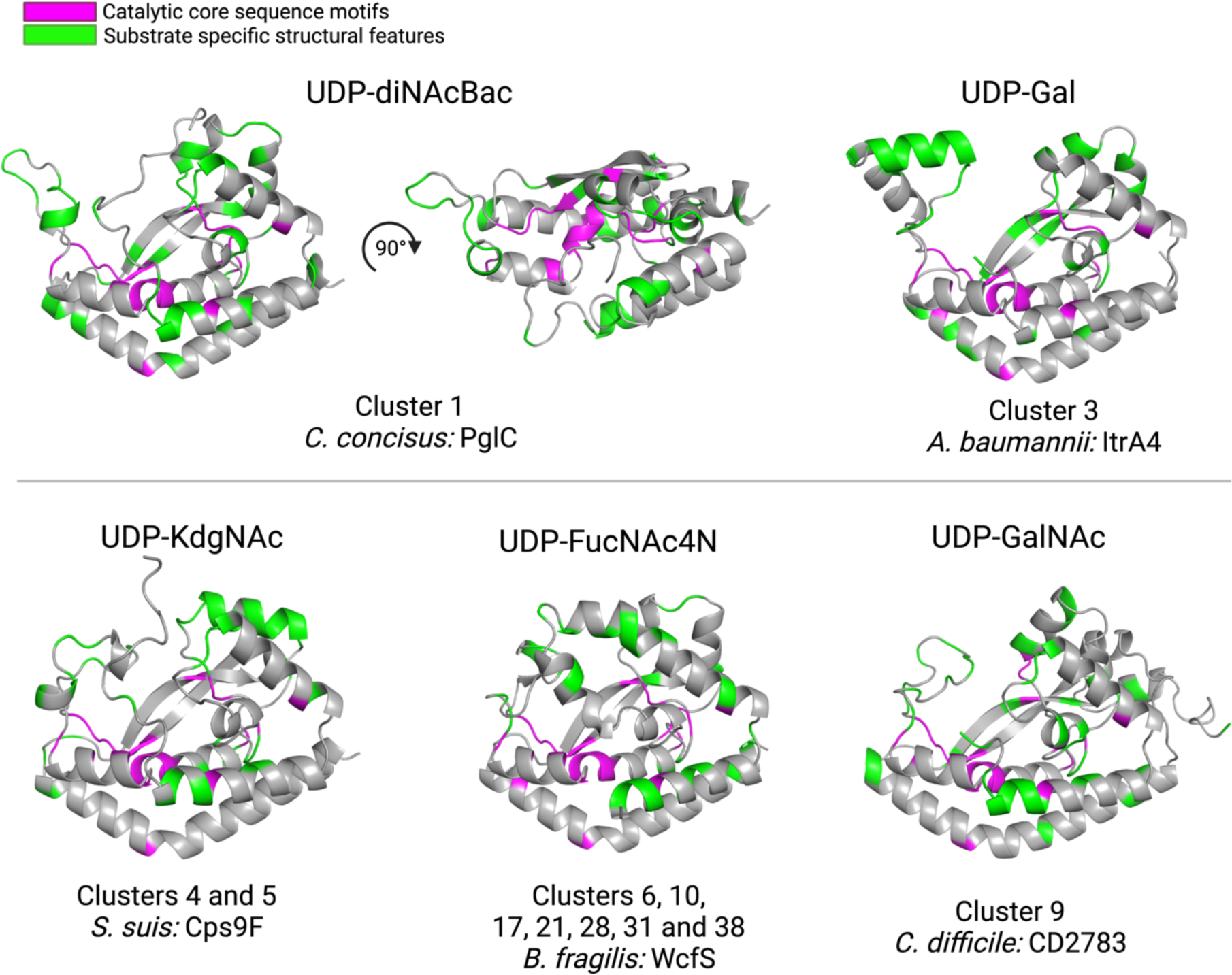
Representative AlphaFold 2 predicted structures of PGTs with different verified UDP-sugar substrates. Catalytic core sequence motifs in magenta are highly conserved across the entire network. Structural motifs in green are enriched in PGT clusters associated with the respective sugar substrates as quantified by 3Di alignment

#### Kinetic Analysis and Inhibitor Screening

The UMP-Glo assay has previously been used for both mechanistic analysis and inhibitor profiling of detergent-solubilized MonoPGTs (10, 19, 36). Therefore, in addition to screening for substrates, we assessed the utility of the CEF-Glo assay as a high-throughput method to assess PGT activity and as a platform to screen SmPGTs against small molecule inhibitors. Kinetic profiles for SmPGT activity were generated using CEF-Glo by quenching reactions at selected time points allowing for determination of linear range conditions (Fig. S10). Once linear range conditions were determined, we introduced selected compounds from an in-house library of nucleoside analogs (36, 37) into the reaction (Fig. 6). Although only a limited number of compounds is assessed as a proof of concept, there is a notable difference in the inhibitory profile of SmPGTs, which allows selection and further analysis of differences in inhibition through concentration-dependent assays (Fig. 6). The best and worst inhibitors of each SmPGT were selected to see if differences in inhibition could be seen in a dose-dependent manner. Indeed, compound 3 (Fig. 6, Fig. S11), a consistently good inhibitor of *C. concisus* and *S. suis*, showed greater inhibition at higher concentrations whereas poorer inhibitors, such as compound 8 (Fig. 6, Fig. S11), showed little improvement in inhibition even at higher concentrations. These distinct inhibitory profiles will now allow for the rationalization of patterns of inhibition across SmPGTs utilizing different substrates. For instance, compounds 6 and 7 (Fig. 6, Fig. S11) are poor inhibitors of the UDP-diNAcBac-utilizing *C. concisus* PglC, but better inhibitors of SmPGTs that use different substrates. In particular, compound 7 (Fig. 6, Fig. S11) is the best inhibitor for the UDP-GalNAc-utilizing *C. difficile* CD2783. Similarly, there are some compounds that consistently inhibit well (compounds 2-4) and inhibit poorly (compound 1). Conclusions from these initial inhibitor screens will enable future expansion and testing of the complete library (ca. 120 nucleoside analogs).

**Figure 6:**
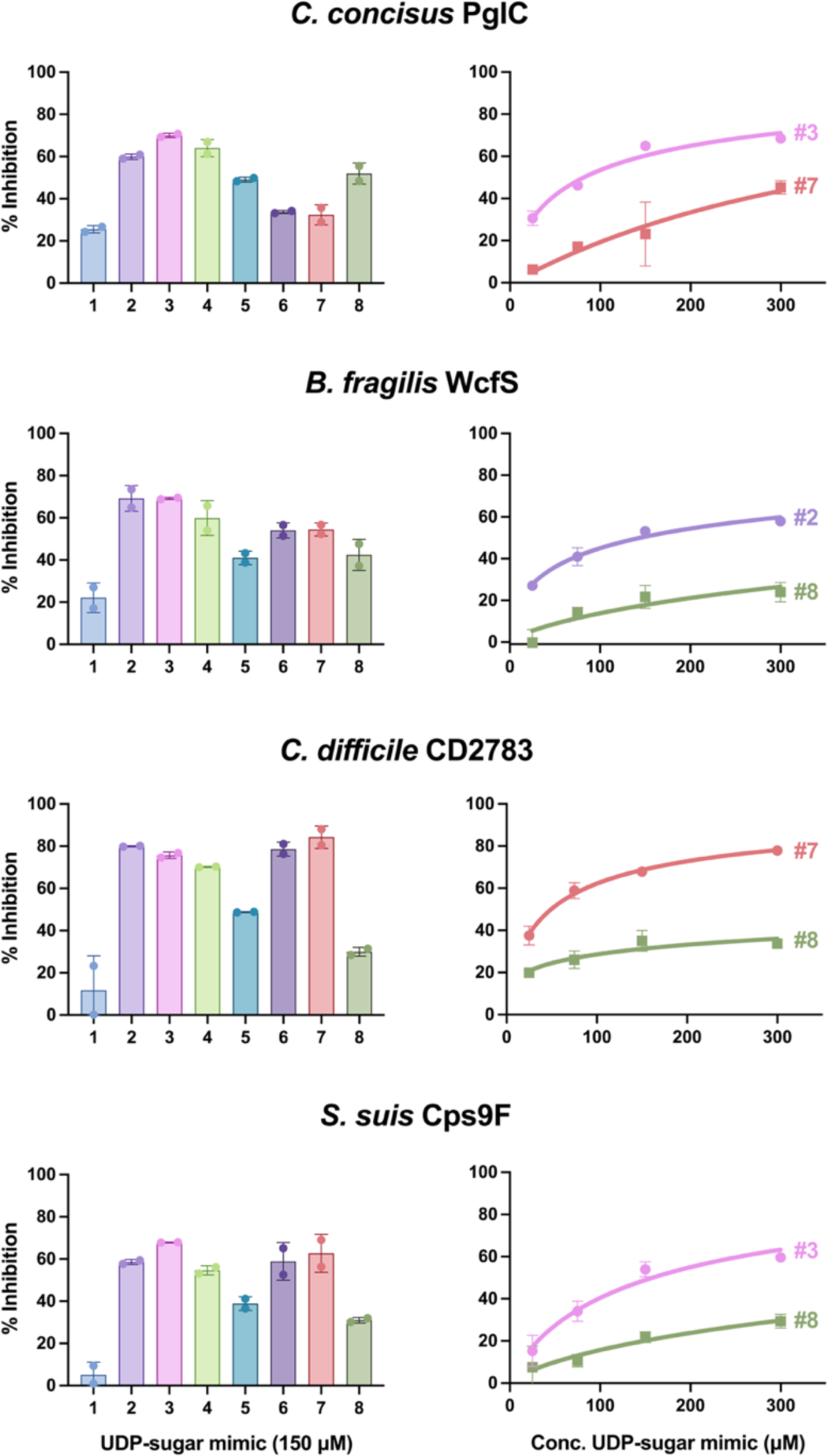
Inhibitor screen and concentration-dependence studies. **Left**: Inhibitor screen assays were run with 150 μM UDP-sugar mimics for a length of time previously determined by linear-range kinetics assays for each SmPGT CEF. Luminescence measurements were recorded in duplicate and observed activity was converted to percent inhibition based on an un-inhibited control. **Right**: The best candidate and a poor candidate were chosen for each SmPGT CEF and used to track concentration dependence over a range of concentrations to confirm inhibition. The same transformation used in the screens were used for the concentration dependence studies.

## Discussion

Expanding the knowledge base of monoPGT substrate specificity to the level of direct biochemical measurement of enzymatic activity provides critical insight into bacterial glycoconjugate biosynthesis. Currently, the functional assignment of enzymes encoded by genes within bacterial biosynthetic operons is typically based on information on native glycoconjugate structures. This approach has long been the cornerstone for PGT and GT function prediction. However, determining glycoconjugate structures requires labor-intensive processes, including isolation and purification from native bacterial sources, followed by sophisticated NMR and/or mass spectrometry-based approaches (Fig. 7). These studies are often complemented by gene knockout experiments, which seek confirmation of function through pathway intermediate build-up. However, such analyses can be misleading in bacterial glycoconjugate pathways as genetic knockout results may be ambiguous when predicting the functions of enzymes that either biosynthesize or use highly-modified carbohydrate building blocks (38). PGT and GT function assignment issues are further compounded by database mis-annotations based solely on primary sequences and predicted structures, a lack of standardized nomenclature for modified bacterial monosaccharides, and the challenge of predicting the beginning and end of repeating units in glycopolymer and glycoconjugate structures *de novo* (39, 40).

**Figure 7:**
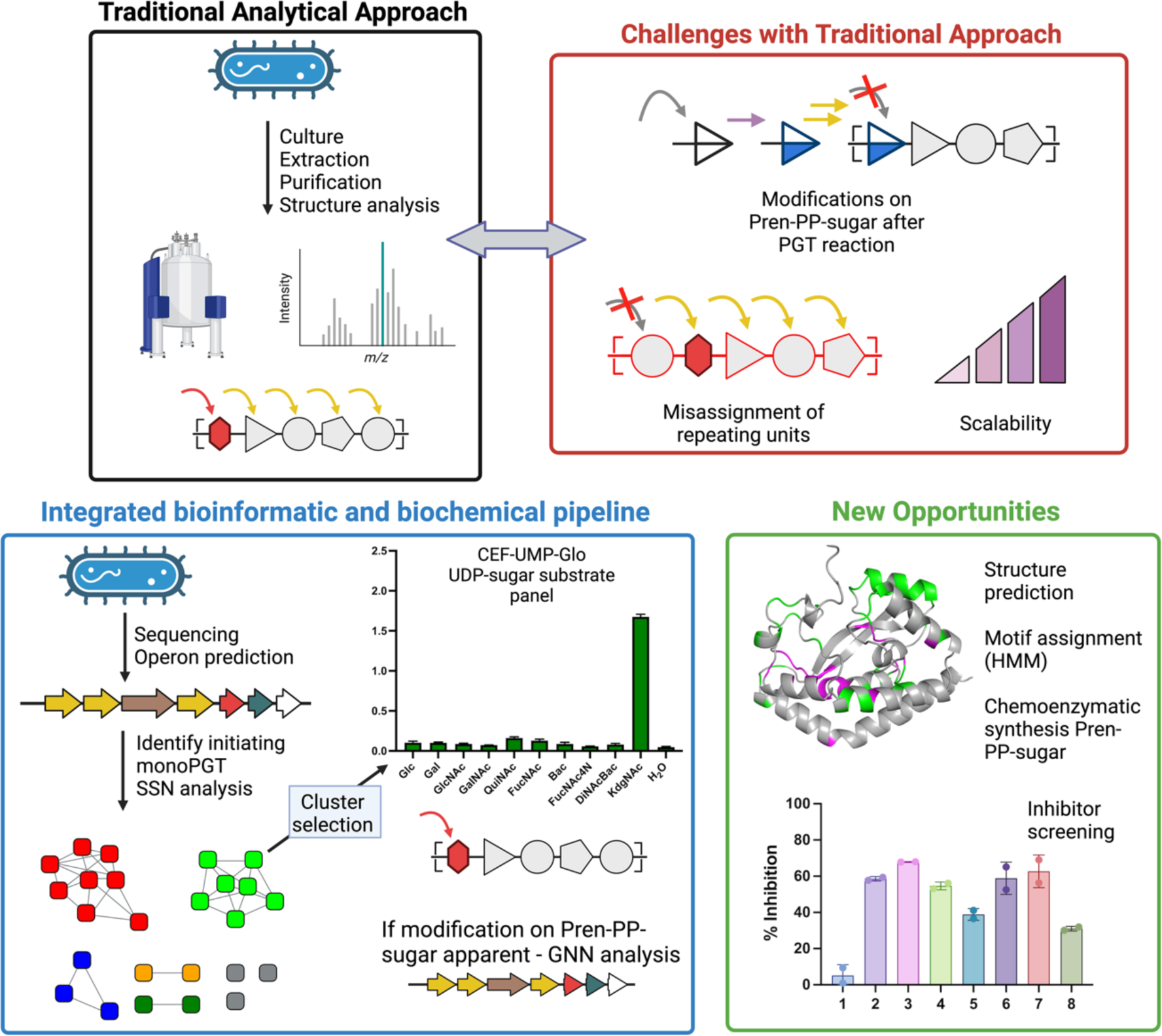
Glycan and glycopolymer structure determination. Traditional methods rely on labor-intensive analytical approaches that have limitations in scale and accuracy of assignment. The integrated bioinformatic and biochemical pipeline affords new information on the superfamily of prokaryotic monoPGTs that initiate glycan biosynthesis and the products of their associated pathway enzymes.

In this work, we couple a bacterial proteome-wide SSN analysis with a targeted luminescence-based biochemical assay compatible with unfractionated CEFs to directly assign and curate the function of the monoPGTs superfamily, which initiates glycan assembly pathways. Knowledge of monoPGT function and substrate specificity will significantly facilitate the study of bacterial glycoconjugates, defining the first carbohydrate added during biosynthesis and informing on the reducing end sugar identity in a broad range of bacterial glycoconjugates. From a practical perspective, knowledge of monoPGT specificity will also provide the acceptor substrate for the following enzyme in each pathway. The isofunctional clusters generated in the current analysis have allowed the prediction of the substrate specificity of nearly 45% of the unique (based on a 95% sequence identity) SmPGT universe (Fig. 3, Table S3), generating valuable information for the field. By uncoupling the PGT assay from the requirement of detergent-solubilized protein, PGT function can be assessed at a much higher throughput and scale than previously possible. We anticipate that future optimization to miniaturize and automate the CEF-Glo assay will achieve greater parity between the scale of bioinformatics analysis and that of functional characterization. In addition, such improvements will facilitate the screening of larger small-molecule libraries, expanding the chemical search space for PGT inhibitors.

Furthermore, as the monoPGT catalytic core domain is highly conserved throughout the superfamily, the current proteome-wide SSN for the SmPGTs can also be exploited to assign substrate specificity for the Bi and Lg monoPGT families. The concepts presented in this study set the stage for the more ambitious goal of predicting the specificity and order of sequential GT-A and GT-B-fold GTs in a glycan assembly production line (41).

### Carbohydrate diversification after PGT activity

Bacterial glycans feature a wide variety of carbohydrate building blocks. Functional prediction of gene products from operons, based on final glycoconjugate structure, can provide insight into PGT and GT function and the identity of cognate NDP-sugar substrates. However, there are a growing number of examples of sugar modification occurring on the UndPP-linked sugar intermediate rather than the UDP-sugar, which can obscure the observed PGT structure-function rationale (42, 43). For example, in *Rhizobium etli* O-antigen biosynthesis, determination of the native LPS structure as containing a reducing-end QuiNAc residue led to an initial prediction of UDP-QuiNAc as the SmPGT (WreU) substrate (Cluster 5). However, WreU did not show PGT activity against the proposed nucleotide substrate. Additionally, the dehydrogenase enzyme predicted to generate UDP-QuiNAc in the operon (WreQ) displayed only marginal activity with UDP-KdgNAc, which suggests this proposed transformation is unlikely to be physiologically relevant (44). It was then demonstrated that the C4-keto biosynthetic precursor, UDP-KdgNAc is the actual PGT substrate, affording UndPP-KdgNAc, which is then converted to UndPP-QuiNAc by the dehydrogenase WreQ (43). The CEF-Glo assay results confirm this sequence of events and further illuminate the function of a large cluster of SmPGTs with specificity for UDP-KdgNAc in the SSN.

A similar logic is at play in SmPGTs from other microorganisms that clearly cluster as utilizing UDP-4-keto-sugar substrates. For example, capsule biosynthesis in the *Vibrio vulnificus* M06-24 pathway is initiated by the SmPGT WbfU (45), (Cluster 7), which we show can transfer the phospho-sugar from UDP-KdgNAc to UndP, to form UndPP-KdgNAc (Fig. 4). However, the reducing-end sugar in the *V. vulnificus* capsule repeating unit is not KdgNAc (46), but instead, a sugar (L-QuiNAc) whose biosynthesis is thought to derive from a UDP-4-keto-sugar similar to UDP-KdgNAc (47). In light of this data, we hypothesize that the biosynthetic steps leading to L-QuiNAc may occur on the UndPP-linked sugar. Supporting this, several other PGTs in Cluster 7 are present in bacteria generating glycoconjugates with highly-modified monosaccharide reducing ends (48-50), with reported soluble biosynthesis from the above UDP-4-keto-sugar (51, 52). Although requiring further investigation, this demonstrates how the current analysis allows us to rationalize monoPGT structure-function assignments which appear as incompatible with the isofunctional clustering.

In some instances, we do not observe activity for the predicted UDP-sugar in the screen even though the assignment appears well-substantiated (Fig. S8). For example, the SmPGT, Cap5M, in the *S. aureus* capsular polysaccharide biosynthesis operon is clustered at E-value 61 with PGTs that act on UDP-KdgNAc. However, under our assay conditions, Cap5M does not catalyze the transfer of this phospho-sugar. In this case, the lack of activity may be the result of regulation at the PGT step; for example activation by tyrosine phosphorylation has been reported (53). *R. leguminosarum* PssA also failed to show activity in the CEF-Glo assay but transfer of phospho-[^3^H]Glc was observed by using radiolabeled substrate in a liquid-liquid extraction assay (Fig. S12). This highlights the possibility that CEF-Glo may be incompatible with some candidate PGTs, due to assay sensitivity or because of molecular interactions in the crude membrane fraction due to the prevalence of metabolic enzymes that utilize common UDP-sugars. Other negative results for SmPGTs may represent areas where the, necessarily-limited, UDP-sugar panel does not include the cognate NDP-substrate of the target SmPGT, as could be the case for the *V. parahaemolyticus* SypR target.

### Expanding the nucleoside diphosphate-sugar panel

Currently, the range of UDP-sugar substrates that are available limits the biochemical analysis of SmPGTs. However, as more structure/function relationships are advanced and as we further the analysis of Genome Neighborhood Networks and Diagrams (GNNs and GNDs) (21, 54, 55) to provide insight into SmPGT-initiated glycoconjugate pathways, new UDP-sugar targets for chemical or chemoenzymatic synthesis will emerge and we anticipate that the substrate specificity of additional clusters from the SSN will continue to be elucidated.

As an example, Cluster 24 contains a SmPGT (Uniprot: Q814Z7) from *Bacillus cereus* (strain ATCC 14579). This bacterium is reported to biosynthesize an unusual 2-oxoglutarate modified UDP-amino sugar (UDP-Yelosamine) using enzymes directly upstream of the SmPGT in the biosynthetic operon (Fig. S13) (56).The GND of cluster 24 shows that the ATP-grasp domain protein responsible for transfer of the oxoglutarate moiety, as well as other sugar-modifying enzymes responsible for generation of the corresponding UDP-sugar intermediates, are largely present in the vicinity of the PGTs in the cluster. This allows us to predict that the cluster may represent SmPGTs with UDP-Yelosamine substrate. This observation now provides an opportunity to investigate and potentially assign a monoPGT cluster with unknown specificity.

### An evolutionary perspective of the MonoPGT SSN

The substrate scope of SmPGTs reveals an interesting logic in the evolution of carbohydrate utilization in bacterial glycoconjugates. SmPGTs in clusters that separate between E-value 61 and E-value 71, mainly to act on simple and abundant UDP-Glc and UDP-Gal substrates, whereas those differentiating at more stringent E-values use modified UDP-sugars such as UDP-diNAcBac, UDP-KdgNAc and UDP-FucNAc4N (Fig. 3). The generation of these modified UDP-sugar substrates requires a keto intermediate, and the ability of the carbonyl group to afford a site for further modification provides provides a strategy for diversifying glycoconjugate pathways.

Bacterial pathogens and symbionts have converging evolutionary pressures for generation of highly-modified extracellular polysaccharides; pathogens increase diversity to avoid recognition by the immune response of their host, whilst symbionts increase diversity to enable more robust interactions with their co-evolving organisms (57). From this lens, it is interesting to see that both highly pathogenic (*S. aureus*, *B. fragilis* and *Streptococcus sp*) and symbiotic (*R. etli*, *A. fischerii* and *Photobacterium sp*) bacteria are over-represented within clusters of SmPGTs that use modified substrates. This is also the case for bacteria that are known opportunistic pathogens, able to live in symbiosis for extended periods before becoming pathogenic if the host is perturbed (*F. nucleatum*, *S. pneumoniae and Burkholderia sp*). By rationalizing these observations to the use of highly modified UDP-sugar substrates, either by the monoPGT or the later GTs we propose an evolutionary niche for the keto-sugar intermediates and derivatives as well as their sugar-appending pairs. In this way, the keto-sugar-producing dehydrogenase, and to some extent the co-evolving monoPGT, can be seen as chemical gatekeepers of glycoconjugate diversity.

### Inhibitor opportunities and screening

In parallel, the CEF-Glo assay platform can be used to directly screen inhibitor libraries on the monoPGT family in a high-throughput modality, facilitating the generation of novel inhibitors as leads for antibiotics and antivirulence agents for this functionally important family. From our preliminary screening data, it is clear that distinct inhibition profiles that differ by chemotype of inhibitor exist across SmPGTs of varying substrate specificity. These results were confirmed through concentration-dependence studies showing a dose-dependent response. This opens the possibility of leveraging this variation in chemotype to differentially inhibit one PGT over another even within the same organism. Conversely, these inhibitor profiles could be leveraged in the design of “broad-spectrum” inhibitors to act on monoPGTs in different bacteria. The assay itself will facilitate the screening of more complete nucleoside-analog libraries against monoPGTs found in pathogenic bacteria, aiming to identify lead compounds for subsequent optimization.

### Summary

Here, we have applied sequence similarity network (SSN) analysis with a luminescence-based activity assay, to probe the extensive family of bacterial monoPGTs. The open availability of the proteome-wide monoPGT data will facilitate the study of bacterial glycoconjugate pathways and will be amenable to expansion and collaboration with researchers in the field through a curated website. Our analysis reveals substrate-specificity at a far more significant scale than previously achievable and builds a foundation for future exploratory campaigns. By providing both the means for biochemical investigation of PGTs and bioinformatic prediction and rationalization of enzyme activity we have begun to lift the veil on the glycan-based strategies employed by bacteria for interaction with each other, their hosts, and with their environment.

## Materials and Methods

### Protein Expression

Gene fragments for each SmPGT indicated in Table 1 apart from WbfU were codon optimized for *E. coli* expression and ordered with 3’ and 5’ overlaps to pe-SUMOpro vector, generating an N-terminal 6-His SUMO construct (His-SUMO-PGT) or C-terminal His_6_ tagged (PGT-His_6_) construct (*C. concisus* PglC and *C. botulinum* SmPGT). Fragments were assembled using the HiFi Assembly kit (New England Biolabs). Plasmid encoding WbfU with N-terminal 6-His NusA tag in the pET-50b vector was generously donated by J. Troutman. Sequences were verified with Sanger sequencing. Plasmids were transformed into *E. coli* cells and grown in autoinduction media in the presence of appropriate antibiotic at 37 °C. Once OD_600_ reached 1.5, the temperature was adjusted to 18 °C, and BL21-AI cells were induced with 1g L-arabinose. Cells were harvested via centrifugation after 18 h of incubation and stored at -80°C.

#### CEF Isolation

Each step of isolation was performed on ice. Frozen pellets were dispersed manually and resuspended using 4 mL per g of pellet of Buffer A (50 mM HEPES pH 7.5, 150 mM NaCl). Contents were transferred to a sonication cup and supplemented with 2 mM MgCl_2_, 0.06 mg/mL lysozyme (RPI), and 0.5 mg/mL DNase I (Millipore Sigma) and left to incubate for 30 min. Cells were lysed using sonication (2x, 50% amplitude, 1s on 2 s off), and CEF was isolated by differential centrifugation (Ti-45 rotor, 9,000g 45 min, reserve supernatant, 140,000g 65 min, decant supernatant). The resulting pellet was homogenized in 25 mL of buffer A and 1 mL aliquots were reserved for assay and flash frozen in liquid N_2_ for storage at -80 °C.

### Chemoenzymatic synthesis of UDP-sugars

UDP-diNAcBac and UDP-Bac were prepared as described previously (30).

#### UDP-KdgNAc generation

UDP-KdgNAc was prepared based on the previously published protocol with minor modifications (30). The plasmid of the *Campylobacter jejuni* 11168 PglF gene with truncated N-terminal transmembrane domains (residues 1-130) and N-terminal GST-tag in pGEX-4T-2 vector was transformed into BL21 cells grown in MDG media supplemented with carbenicillin at 37 °C for 18 h with protein production being induced with 1 mM IPTG. Cells were harvested and stored at -80 °C.

For GST-PglF_Δ1-130_ immobilization, cells were lysed in 40 mL Buffer A, supplemented with 25 mg lysozyme, 25 μL DNAse I and 40 μL protease inhibitor cocktail (EMD Millipore 539134) before sonication as described above. Clarified lysate was isolated by centrifugation of homogenate (Ti-45 rotor, 35,000 rpm 1h). Lysate was incubated with 4 mL Glutathione agarose resin (PierceTM), pre-equilibrated with Buffer A in the presence of 1 mM of NAD^+^ (Sigma N0632) for 2 h at 4 °C. Resin was transferred to a column and the excess clarified lysate was flowed through the column via gravity. The column was washed with 8-column volumes (CV) of ice-cold Buffer A to remove excess protein. The reaction was initiated by addition of 25 mg of UDP-GlcNAc (Sigma U4375) dissolved in 1 CV of Buffer A with 1 mM NAD^+^ to the resin. The reaction was incubated at room temperature with gentle rotation for 48 h. The flow-through containing the product was collected and the resin was washed with 2 CV of reaction buffer. The unwanted soluble protein in the flow-through and wash fractions were removed by heating the solution at 60 °C for 1 h, followed by centrifugation at 3,200 x g for 30 min. The crude product was purified by anion-exchange HPLC equipped with semi-preparative column with guard column (Column: Thermo Scientific Dionex CarboPac^TM^ PA1, 9 x 250 mm, Guard Column: Dionex CarboPacTM PA1 9 x 50 mm). A mobile phase of, A: ddH_2_O and B: 1 M NH_4_HCO_3_, with gradient 3% to 20% B over 10 min, 20% to 35% B over 30 min, 35% to 50% B over 15 min and 50% to 100% B over 5 min was used (Gradient A). UDP-KdgNAc peak was isolated by comparison of retention time to an authentic standard. The combined fractions from HPLC elutions containing the desired product were collected, lyophilized (37% yield, 9.4 mg) and characterized by ^1^H NMR and ESI (-) LRMS (Fig. S4).

#### Reduction of UDP-KdgNAc to UDP-QuiNAc and UDP-FucNAc

Based on a similar procedure with the corresponding TDP-sugar (31), we obtained the C4’ hydroxyl epimers - UDP-QuiNAc and UDP-FucNAc by NaBH_4_ reduction of UDP-KdgNAc. UDP-KdgNAc (1.5 mg, 2.5 µmol) was transferred to a round bottle flask. Then 9.45 mg NaBH_4_ (Fluka Analytical 71321) suspended in 1 mL of methanol-water (1:1 v/v) was added to the reaction at an equimolar ratio. The reaction was incubated for 2 h at room temperature with gentle stirring, before quenching with 1 mL acetone. Before purification, excess acetone was removed from crude product under nitrogen gas. The reaction product was purified by anion-exchange HPLC (A: ddH_2_O, B: NH_4_HCO_3_) using a gradient of 3% to 22% B over 10 min, 22% to 23% B over 40 min, and 23% to 100% B over 5 min (Gradient B). This gradient was determined through iterative adjustment to be optimal for separation of the stereoisomers but may require optimization for different equipment and to account for concentration variation caused by volatility of the ammonia in buffer B. The eluted peaks were collected, lyophilized (0.18 and 0.06 mg of UDP-QuiNAc and UDP-FucNAc, respectively) and characterized by ^1^H NMR by comparison to published spectra (44, 58) (Fig. S5).

#### WcfR purification

A plasmid encoding His-tagged WcfR aminotransferase was generously provided by J. Troutman. The WcfR sequence was verified by Sanger sequencing, transformed into E. coli BL21 cells and expressed in autoinduction media as described above. Cells were pelleted via centrifugation, and stored at -80 °C.

Cells were resuspended in 50 mM HEPES, pH 8.0, 200 mM NaCl (Buffer B) using 4 mL buffer B per gram of wet cell mass, supplemented with DNase (Roche), MgCl_2_, and lysozyme (RPI) to a final concentration of 0.5 mg/mL, 2 mM, and 0.06 mg/mL respectively. Cells were incubated on ice for 30 min, and then lysed via sonication (2x 50% amplitude, 1 s on 2 s off). Insoluble material was removed via centrifugation (Ti-45 rotor, 42,000 RPM 60 min). The supernatant was filtered and loaded onto a 5 mL HisTrap column (Cytiva) using an Akta FPLC equilibrated with buffer B. Protein was eluted using a linear gradient from 0 – 100% buffer C (50 mM HEPES pH 8.0, 200 mM NaCl, 500 mM imidazole) over 5 column volumes. Peak fractions were pooled, dialyzed overnight against buffer B, aliquoted and stored at -80 °C.

#### WcfR reaction

UDP-FucNAc4N was generated by incubation of 1 mM UDP-KdgNAc with 0.16 mg of purified WcfR, 15 mM L-glutamate, 100 μM of PLP, 50 mM Tris pH 8.0 and 50 mM NaCl at 28 °C for 6 h. WcfR was filtered out with Amicon ultracel 10K centrifuge filters spun at top speed for 15 min. UDP-FucNAc4N was isolated by anion-exchange HPLC using gradient of 3% to 20% B over 10 min, 20% to 60% B over 30 min and 60% to 100% B over 4 min (Gradient C). UDP-FucNAc4N-containing elution-fractions were lyophilized and confirmed to be desired product by ESI (-) LRMS before undergoing CIAP treatment (Fig. S6).

#### CIAP Purification of UDP-sugars

Chemoenzymatically derived UDP-sugars underwent CIAP treatment. This was performed by incubation of UDP-sugar with 0.5 μL of CIAP (5 units) in 50 mM Tris pH 8, 100 mM NaCl and 1 mM MgCl_2_ at 37 °C for 2 h. Protein was removed from the reaction by centrifugal filtration with Amicon Ultracel 10K cutoff at max speed before separation of treated material by anion-exchange HPLC (A: ddH_2_O, B: NH_4_HCO_3_) with gradient C (Fig. S7). UDP-sugar-containing elution fractions were collected, lyophilized and resuspended in ddH_2_O for assays.

#### Substrate Assay

The UMP-Glo-based PGT assay followed the previously published general scheme with minor modification (19). The assay buffer consisted of 0.1% Triton X-100, 50 mM HEPES pH 7.5, 100 mM NaCl, 5 mM MgCl_2_ and CEF concentration was determined by absorbance at 280 nm. For each protein, 200 μM UndP, CEF (0.04 mg/mL final) and assay buffer reaction mix was made up (1:1:7 ratio), 10 μL of which was dispensed for each reaction and left at room temperature for several minutes. Reactions were initiated by addition of 1 μL of 200 μM UDP-sugar or ddH_2_O and left to incubate for 1 h. After incubation, each reaction was quenched with a 1:1 ratio of UMP-Glo reagents (10 μL: 10 μL) directly in a 96-well plate for luminescence measurement by plate reader. Luminescence measurements were converted to [UMP] with a standard curve.

#### Kinetics & Inhibitor Analysis

The reaction mix was generated as described above but scaled up for several time points and for a range of CEF concentrations. Reactions were initiated by adding the appropriate equivalents of cognate 200 μM UDP-sugar. At set time points aliquots from reactions were quenched with a 1:1 ratio of UMP-Glo reagents (10 μL: 10 μL) directly in a 96-well plate.

Once the linear range was determined, the reaction mix was made up in a manner analogous to the substrate screen with final CEF concentration at the expected linear range, 9 μL of reaction mix was aliquoted for each reaction. The nucleoside analog dissolved in DMSO (1 μL) was added to all reactions, except for un-inhibited control where DMSO was added, and left to incubate for several minutes. Reactions were initiated by addition of 1 μL of 200 μM cognate UDP-sugar and left to incubate for a pre-determined linear range time before quenching with UMP-Glo reagents at 1:1 ratio directly in a 96-well plate followed by luminescence measurement.

Concentration-dependence studies were carried out in a similar fashion to the inhibitor screen. The best and worst nucleoside analogs were chosen based on the results of the screen. Dilutions of compounds were made to give final concentrations of 25 μM, 75 μM, 150 μM, and 300 μM in the reaction. Reaction mix was aliquoted for each reaction, 1 μL of analog at its given dilution or 1 μL of DMSO was left to incubate with the mixture for several minutes. Reactions were initiated by addition of 1 μL of 200 μM cognate UDP-sugar. Reactions were left to incubate for the pre-determined linear range time and quenched with UMP-Glo reagents at a 1:1 ratio directly in a 96-well plate. This was repeated for each inhibitor.

### SSN Generation

The Uniprot protein database was searched for the IPR ID: PF02397 (Bac-transferase domain) exclusive to the monoPGT family, sequences were pooled and filtered to include only those larger than 150 AA and smaller than 250 AA, consistent with the SmPGT size cut-off. The generation of an SSN was performed in a routine manner using the EFI-EST server (21). Briefly, sequences were inputted in FASTA format, and an initial arbitrary E-value cut-off 1×10^-5^ was set for all-to-all alignment, fragments were excluded, and alignment was limited to the IPR ID: PF02397. After initial alignment was complete, E-value cut-off was adjusted incrementally from E-40 to E-100, generating a series of networks that were taken forward for closeness analysis.

#### Closeness Analysis

To reduce computational time, the Closeness Analysis script was adjusted so that it only generates whole network closeness and not also closeness by cluster. Further, Closeness analysis was performed on 75% collapsed repnode networks, meaning that sequences with greater than 75% identity were collapsed into a single node, minimizing number of edges and therefore computational strain. Closeness centrality was calculated for all networks generated in previous step and plotted (Fig. S1).

#### Dendrogram Generation

Mapping tables generated by EFI-EST, representing the cluster position of each node in the generated networks were transformed to represent the cluster position at E-values 40-71 of a representative sequence for each cluster in the E-71 network. The resulting database was used to write a dendrogram for the largest 50 clusters in the E-71 network in Newick format manually. The dendrogram was visualized using Microreact with Leaf nodes coloring according to E-value 71 cluster coloring (59).

#### 3Di Analysis

The entire network was aligned in sequence space using the Clustal omega algorithm. Catalytic core sequence motifs were identified by selecting residues with the highest Shenkin divergence conservation score (60). Structure alignments were performed with no filtering and default parameters using the Foldseek tools for 3Di encoding and alignment from the Foldmason github repository (35, 61). Of the 32,467 sequences in the network, all but 4,585 had predicted structures in the AlphaFold database. These remaining sequences were folded using ColabFold (34, 62). Dissimilarity was calculated as the column-wise Euclidean distance weighted by the Foldseek 3Di substitution matrix and quantified for both substrate-specific alignments and the network as a whole (Fig. S9). Substrate-specific features were identified as having a 40% or greater dissimilarity in the network alignment than in the substrate group and specific clusters compared for the E-71 alignment are given (Fig. S9).

## CRediT authorship contribution statement

TD, GJD, KNA and BI designed the overall strategy and the experimental approach; TD, GJD and KNA carried out the SSN and GNN analyses; TD, RPS, CAA and SG carried out protein expression, UDP-sugar chemoenzymatic synthesis and CEF Glo analysis; TD, RPS, and CAA carried out inhibition studies; HRH performed 3Di analysis; TD, GJD, and BI wrote the manuscript. All authors reviewed and edited the manuscript.

## Data Availability

The sequence similarity network (SSN) and Genome Neighborhood Network (GNN) files have been deposited in Mendeley (DOI: 10.17632/yg333b6j3b.1). All other data can be found either within the manuscript or in the Supplementary Information (SI).

## Acknowledgements

We thank Prof. Jay Troutman for providing plasmids and for valuable discussions. Figures 1 and 7 were made in BioRender.com.

