## Supporting Information for "Proteome-Wide Bioinformatic Annotation and Functional Validation of the Monotopic Phosphoglycosyl Transferase Superfamily"

[c]Current address Biogen, 225 Binney Street, Cambridge MA 02139, USA

| | | |
|---|---|---|
| Theo Durand | 0009-0004-8131-581X | Dept. of Biology, MIT, Cambridge MA 02139, USA. Dept. of Life Sciences, ICL, London SW7 2AZ, UK |
| Greg J. Dodge | 0000-0002-6555-8350 | Biogen, 225 Binney Street, Cambridge MA 02139, USA |
| Roxanne P. Siuda | 0000-0001-7387-8429 | Dept. of Chemistry, BU and Dept. of Pharmacology, Physiology and Biophysics, BUSM |
| Hugh R. Higinbotham | 0000-0002-9418-7958 | Dept. of Biology and Dept. of Physics, MIT |
| Christine A. Arbour | 0000-0001-6056-296X | Dept. of Biology and Dept. of Chemistry, MIT |
| Soumi Ghosh | 0000-0001-6101-0147 | Dept. of Biology, MIT |
| Karen N. Allen | 0000-0001-7296-0551 | Dept. of Chemistry, BU |
| Barbara Imperiali | 0000-0002-5749-7869 | Dept. of Biology and Dept. of Chemistry, MIT |

**Table of Contents**

**Supplementary Tables**

**Table S1:** Comprehensive table of smPGT assignments based on literature precedent

**Table S2:** Common and alternate names of biochemically-modified UDP-sugar substrates for monoPGTs

**Table S3:** The 95% repnode SSN cluster count for assignment of unique SmPGT sequences

**Supplementary Figures**

**Figure S1:** Closeness Analysis of SSNs showing inflection points at E-value 61 and 71

**Figure S2:** SSN Position of SmPGTs presented in the manuscript

**Figure S3:** Expression of the smPGTs in unfractionated Cell Envelope Fractions (CEF) by western blot analysis

**Figure S4:** Characterization of UDP-KdgNAc from chemoenzymatic synthesis

**Figure S5:** Characterization of UDP-FucNAc and UDP-QuiNAc from chemoenzymatic synthesis DP-QuiNAc and UDP-FucNAc chemoenzymatic synthesis

**Figure S6:** Characterization of UDP-FucNAc4N from chemoenzymatic synthesis

**Figure S7:** Calf Intestinal Alkaline Phosphatase (CIAP) treatment for removal of nucleotide contaminants

**Figure S8:** CEF-Glo control data

**Figure S9:** Example Structural Conservation analysis for chosen smPGT

**Figure S10:** Kinetic analysis of selected smPGTs in CEF

**Figure S11:** Radioactivity-based substrate assay with CEF containing overexpressed *R. leguminosarum* PssA

**Figure S12:** Structures of nucleoside analogs used for inhibitor screening

**Figure S13**: UDP-Yelosamine biosynthesis and operon analysis

**Supplementary Methods**

**Typical chemical synthesis scheme of nucleoside analogs**

**Table S1**: Comprehensive table of smPGT assignments based on literature precedent

| Organism | PGT Name | Uniprot | Putative Substrate(s) | E-71 cluster | Glycoconjugate | Reference |
|---|---|---|---|---|---|---|
| *R. etli* | WreU | Q2K1T1 | KdgNAc | 5 | O-antigen/LPS | (1) |
| *F. tularensis schu S4* | WbtB | Q5NEZ2 | KdgNAc/ QuiNAc | 4 | O-antigen/LPS | (2, 3) |
| *S. suis serotype 9* | Cps9F | Q9RG41 | KdgNAc | 4 | CPS9 | (4) |
| *S. pneumoniae serotype 5* | | Q7WVW9 | KdgNAc | 4 | CPS5 | (5) |
| *S. aureus* | Cap5M | P95706 | KdgNAc/ FucNAc | 50 | Type 5 CPS | (6, 7) |
| *S. saprophyticus* | | Q4A111 | KdgNAc | 50 | CPS | (8) |
| *F. nucleatum strain 25586* | | Q8R6F8 | QuiNAc | 132 | O-antigen/LPS | (9) |
| *C. botulinum* | | A0A6B4JHH5 | FucNAc4N | 10 | N/A | |
| *B. fragilis* | WcfS | Q93QV6 | FucNAc4N | 10 | CPSA | (10) |
| *F. nucleatum ATCC 23726* | | D5RFH4 | FucNAc4N | 10 | O-antigen/LPS | (11) |
| *F. nucleatum ATCC 51191* | | F9EN55 | FucNAc4N | 10 | O-antigen/LPS | (12) |
| *F. nucleatum ATCC 10953* | | A5TVH5 | FucNAc4N | 10 | O-antigen/LPS | (13) |
| *F. nucleatum HM-997 or CTI-07* | | A0A829KYS8 | FucNAc4N | 10 | O-antigen/LPS | (14) |
| *F. nucleatum HM-994 or CTI03* | | ERT37135.1 | FucNAc4N | | O-antigen/LPS | (14) |
| *F. nucleatum MJR 7757* | | A0A133NN30 | FucNAc4N | 10 | O-antigen/LPS | (15) |
| *S. sonnei* | | Q3YTB3, Q9S0U8 | FucNAc4N | 17 | Phase I polysaccharide | (16, 17) |
| *P. alcalifaciens* | | A0A346CL59M 9P0X8, M9P183 | FucNAc4N | 17 | O-antigen O22 and O8 | (18-21) |
| *B. bronchiseptica ATCC BAA-588,* | | A0A0R4J6E5 | FucNAc4N | 17 | Endotoxin/LPS | (22, 23) |
| *S. mitis strain B6* | | D3H7E6 | FucNAc4N | 6 | Cell wall PS/LTA IV | (24, 25) |
| *S. pneumoniae serotype 4* | | A0A0H2URP3 | FucNAc4N | 6 | LTA IV | (26, 27) |
| *S. suis serotype 7* | Cps7F | Q9RFX2 | FucNAc4N | 6 | LTA IV | (28) |
| *S. pneumoniae serotype 1* | | A0A0H2ZRD5 | FucNAc4N | 6 | LTA IV | (29) |
| *H. parainfluenzae T3T1* | | E1W1Z5 | FucNAc4N | | LPS | (30) |
| *H. parainfluenzae strain 20* | | R9WQP1 | FucNAc4N | 10 | LPS | (31) |
| *B. fragilis strain 638R* | | E1WLL6 | FucNAc4N | 21 | PS1 | (32, 33) |
| *R. meliloti* | ExoY | Q02731 | Gal | 3 | Succinoglycan | (34) |
| *R. fredii* | ExoY2 | G9AFL7 | Gal | 3 | LPS | (35) |
| *M. japonicum* | | *Q98C89* | Gal | 3 | LPS | (35) |

| | | | | | | |
|---|---|---|---|---|---|---|
| *A. baumannii B8300* | ItrA4 | A0A2I8CW33 | Gal | 3 | CPS | (36) |
| *A. baumannii B850* | ItrA4 | A0A7S8F8L0 | Gal | 3 | CPS | (37) |
| *L. rhamnosus GG* | EpsE | C1J9J2, A0A809NCL8 | Gal | 14 | EPS | (38) |
| *R. leguminsarum* | PssA | Q52856 | Glc | 18 | EPS | (35) |
| *R. legum. Bv trifolii* | PssA | B2Z9S5/A0A9X 5D0X6 | Glc | 18 | EPS | (35) |
| *L. johnsonii FI9785* | EpsE | D0R4M7 | Glc | 2 | EPS | (39) |
| *V. parahaemo-lyticus serotype O3:K6* | SypR | Q87PP3 | Unknown | 20 | Structural symbiosis PS | |
| *C. difficile* | CD278 3 | Q183M0 | GalNAc | 9 | PS-II | (40, 41) |
| *V. vulnificus M06-24* | WbfU | A0A4Q7IER7 | D-KdgNAc/ L-KdgNAc | 7 | CPS | (42) |
| *A. fischeri* | | Q5E8F5 | D-KdgNAc/ L-KdgNAc | 7 | O-antigen/LPS | (43) |

**Table S2:** Common and alternate names of biochemically-modified UDP-sugar substrates for monoPGTs

| Sugar, name used in this work | Alternative names |
|---|---|
| **KdgNAc** | 2-acetamido-2,6-dideoxy-D-xylose-hexos-4-ulose (**Sug**) (4, 44)<br>2-acetamido-2,6-dideoxy-D-xylo-4-hexulose (**ADHexu**) (3)<br>4-keto sugar (45) |
| **FucNAc4N** | 2-acetamido-4-amino-2,4,6-trideoxy-β-galactopyranoside (14)<br>4-*N*-D-FucNAc<br>2-acetamido-4-amino-2,4,6-trideoxy-D-galactose (16)<br>D-FucNAc4N: 2-acetamido-4-amino-2,4-dideoxy-D-fucose (46)<br>α-D-Fuc*p*NAc4NR (R is partial acetylation of the sugar)<br>2,4-diamino-2,4,6-trideoxydeoxy-D-galactose (12)<br>4-amino-D-FucNAc (47)<br>2-acetamido-4-amino-2,4,6-trideoxy-D-galactose (**AATGal**) (48)<br>2-acetamido-4-amino-6-deoxygalactopyranose (**AADGal**) (10) |
| **diNAcBac** | *N,N′*-diacetylbacillosamine<br>2,4-diacetamido-2,4,6-trideoxy-D-glucose (49)<br>2,4-diNAc: 2,4-diacetamido-2,4,6-trideoxy-α-D-glucopyranose (50)<br>di-*N*-acetyl D-bacillosamine (**D-Bac**) (51)<br>2,4-diamino-2,4,6-trideoxyhexoses (**DATDH**) (52)<br>2,4-diacetamido-2,4,6-trideoxy-D-glucose (**QuiNAc4NAc**) (53)<br>2-acetamido-4-acylamino-2,4,6-trideoxy-D-glucose (**D-QuiNAc4NR**) (54) |

**Table S3:** The 95% representative node (repnode) SSN cluster count for assignment of unique SmPGT sequences. The percent of SmPGT sequence space predicted based on our assignments is: whole network node number divided by the number of nodes in the assigned cluster = 44.3%

| Clusters assigned at 95% repnode | Number of Nodes |
|---|---|
| Whole network | 24677 |
| Cluster 1 | 4514 |
| Cluster 2 | 1373 |
| Cluster 3 | 645 |
| Cluster 4 | 753 |
| Cluster 5 | 705 |
| Cluster 6 | 367 |
| Cluster 7 | 601 |
| Cluster 8 | 477 |
| Cluster 9 | 295 |
| Cluster 10 | 367 |
| Cluster 17 | 242 |
| Cluster 18 | 138 |
| Cluster 21 | 175 |
| Cluster 28 | 105 |
| Cluster 31 | 101 |
| Cluster 38 | 76 |

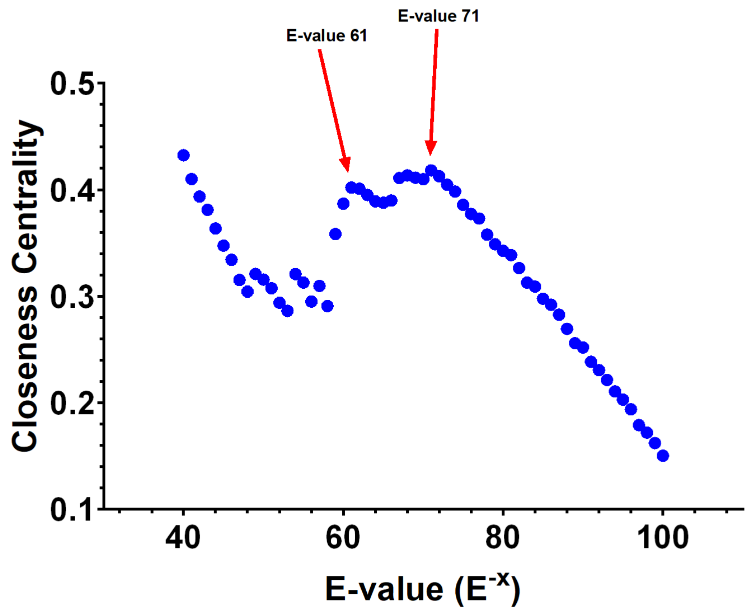


**Figure S1**: Closeness Analysis of SSN showing inflection points at E-value 61 and 71. Closeness centrality was calculated for 75% repnode networks (sequences above a 75% identity collapsed into single nodes) across E-values.

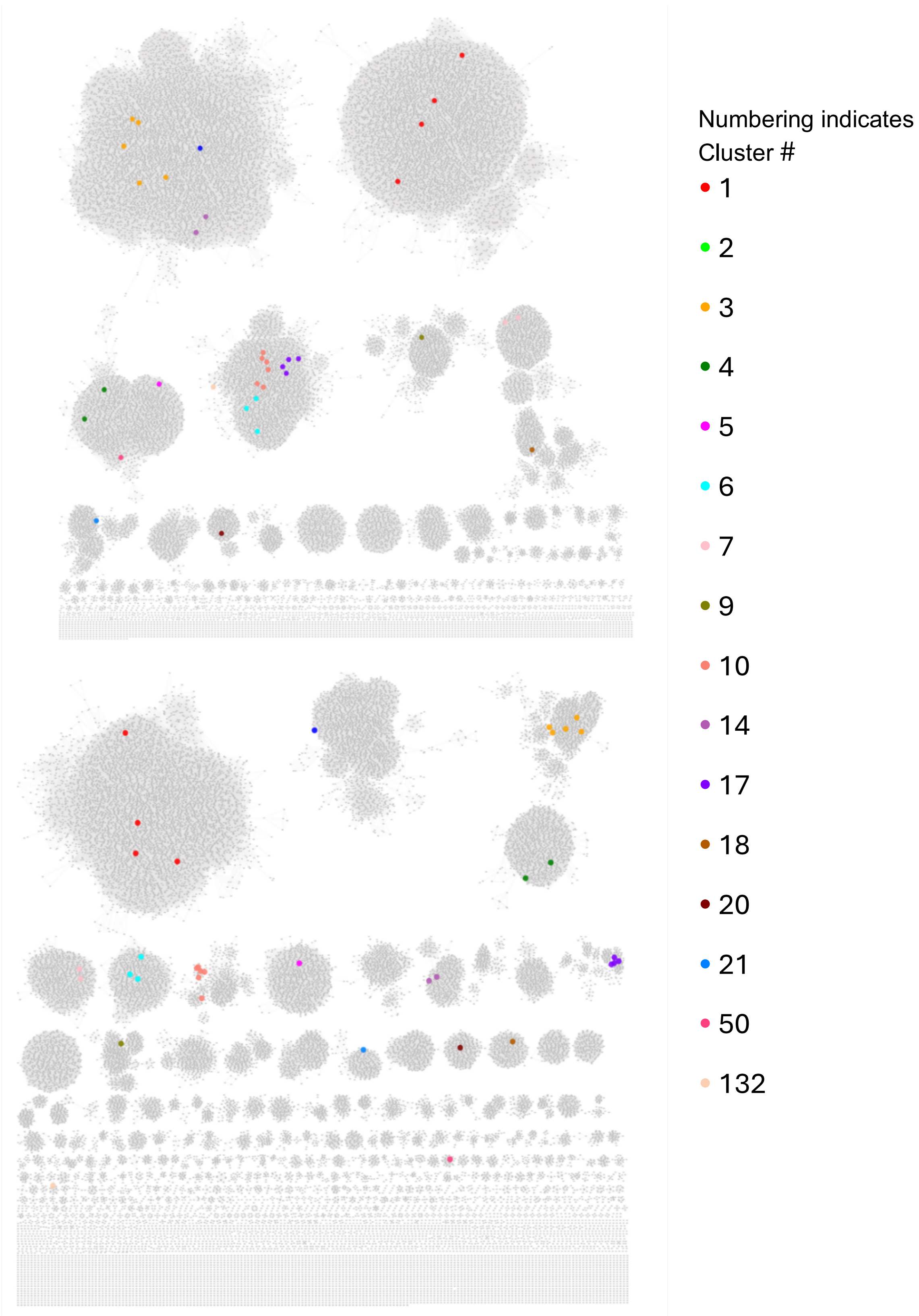


**Figure S2**: SSN Positions of SmPGTs presented in the manuscript. **Top**: E-61 network, **Bottom**: E-71 network. SmPGTs with putative assignment based on Table 1 and S2 are highlighted and color-coded in the E-61 and E-71 networks.

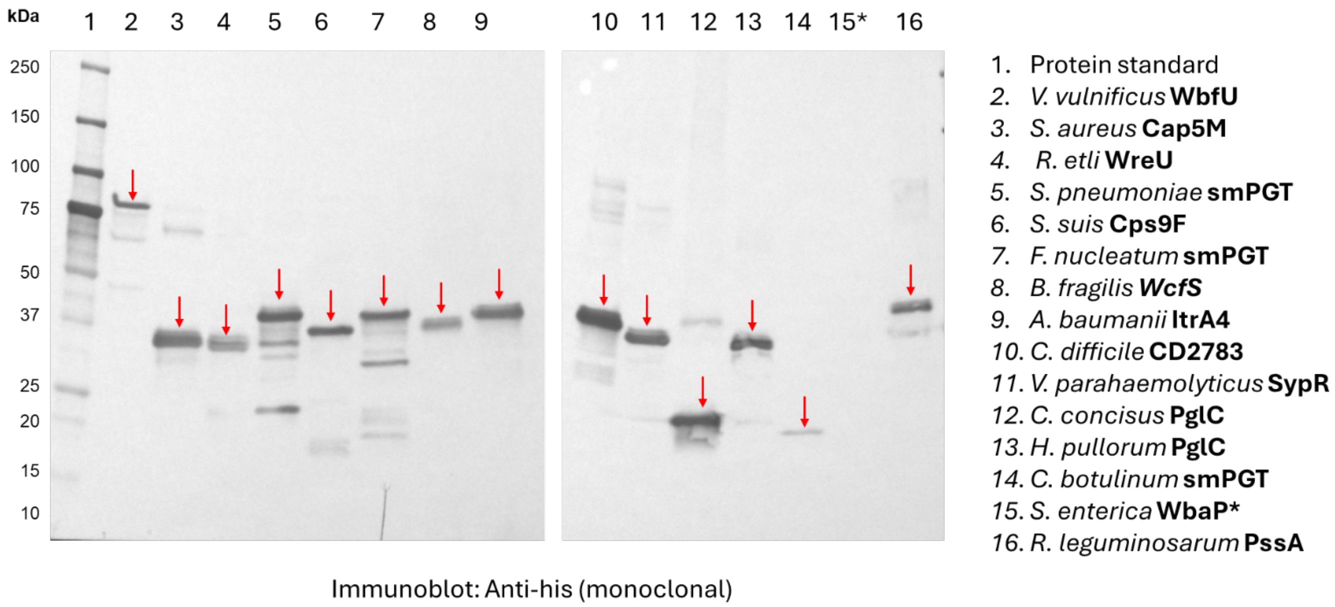


**Figure S3**: Expression of the SmPGTs in unfractionated cell envelope fractions (CEF) by western blot analysis: The CEF of SmPGTs (10 µg total protein, based on 280 nm measurement) were separated by gel electrophoresis, transferred to a nitrocellulose membrane, and probed with monoclonal anti-his antibody (LifeTein). Red arrows indicate the presence of protein at expected molecular weights. **S. enterica* WbaP was not observed on the western blot as it was expressed with a twin Strep-tag in the CEF.

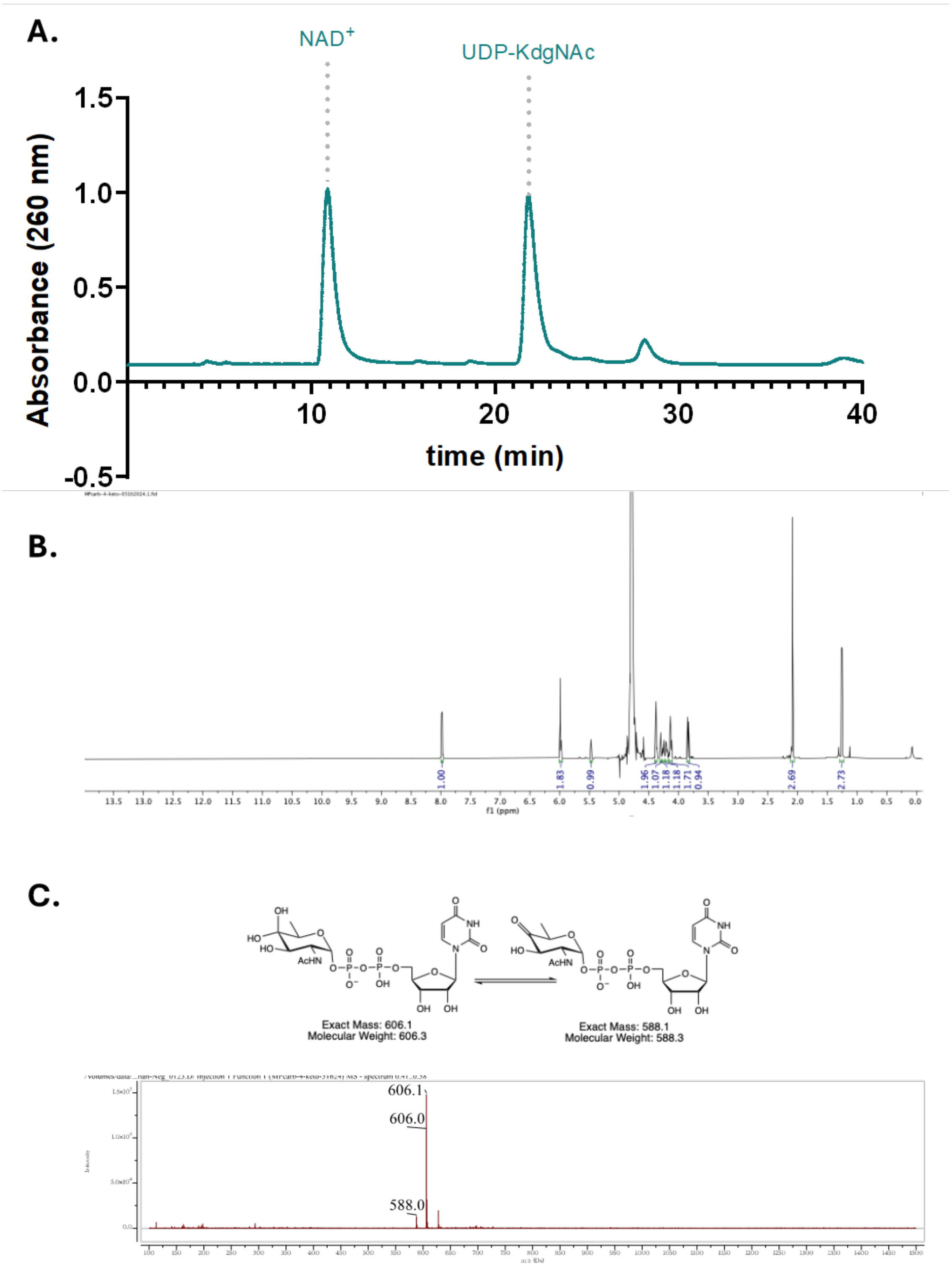


**Figure S4**: Characterization of UDP-KdgNAc from chemoenzymatic synthesis. **A**. A 16 µL injection of 20mM UDP-KdgNAc reaction material using anion-exchange HPLC with gradient A. The first peak is hypothesized to be $NAD^+$ based on equimolar addition of $NAD^+$ and UDP-KdgNAc in the reaction mixture. UDP-KdgNAc was isolated over multiple rounds of purification and elution fractions were lyophilized. **B**. $^1H$ NMR spectrum (600 MHz, $D_2O$) of 1.5 mg of lyophilized elution fractions, confirming identity of the isolated peak as UDP-KdgNAc based on comparison with published spectra (55). **C.** LRMS ESI(-) of lyophilized elutions and exact molecular weight predictions of the hydrated (left) and un-hydrated (right) UDP-KdgNAc species which exist in equilibrium in solution. LRMS ESI(-) confirms the presence of both species in agreement with NMR.

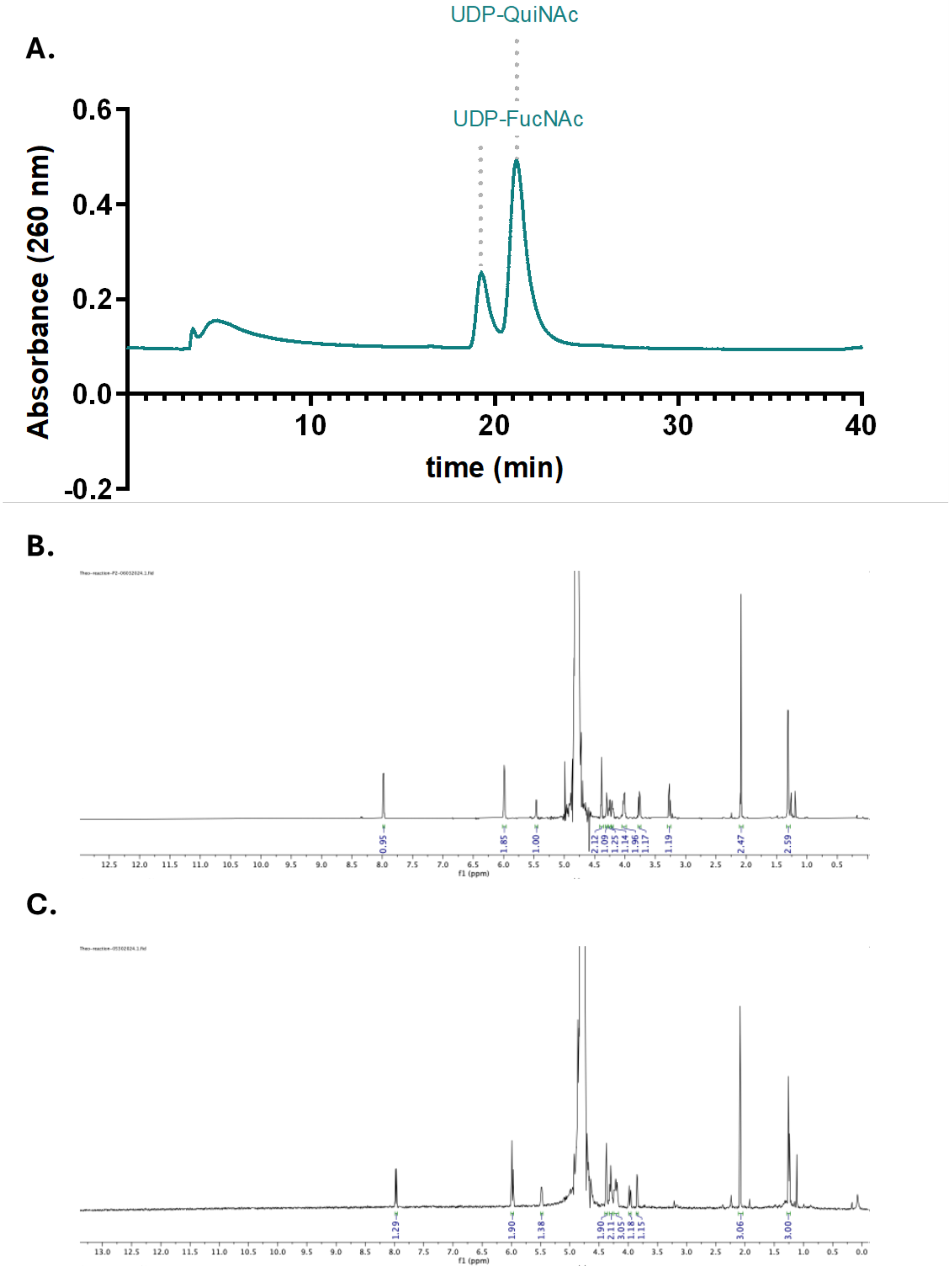


**Figure S5**: Characterization of UDP-FucNAc and UDP-QuiNAc from chemoenzymatic synthesis. **A**. A 400 µL injection of UDP-QuiNAc/UDP-FucNAc reduction reaction using anion-exchange HPLC with gradient B. The broad peak at retention time ~5 min is hypothesized to be residual acetone. The major peaks between 18-23 min were individually isolated over several rounds of purification and lyophilized for analysis generating 0.06 mg of peak 1 and 0.18 mg of peak 2. **B**. $^{1}$H NMR spectrum (600 MHz, $D_2O$) of 0.18 mg of lyophilized peak 2 which was compared to published spectra to identify as UDP-QuiNAc (55). **C**. $^{1}$H NMR spectrum (600 MHz, $D_2O$) of 0.06 mg of lyophilized peak 1 which was compared to published spectra to identify as UDP-FucNAc (56).

**A.**

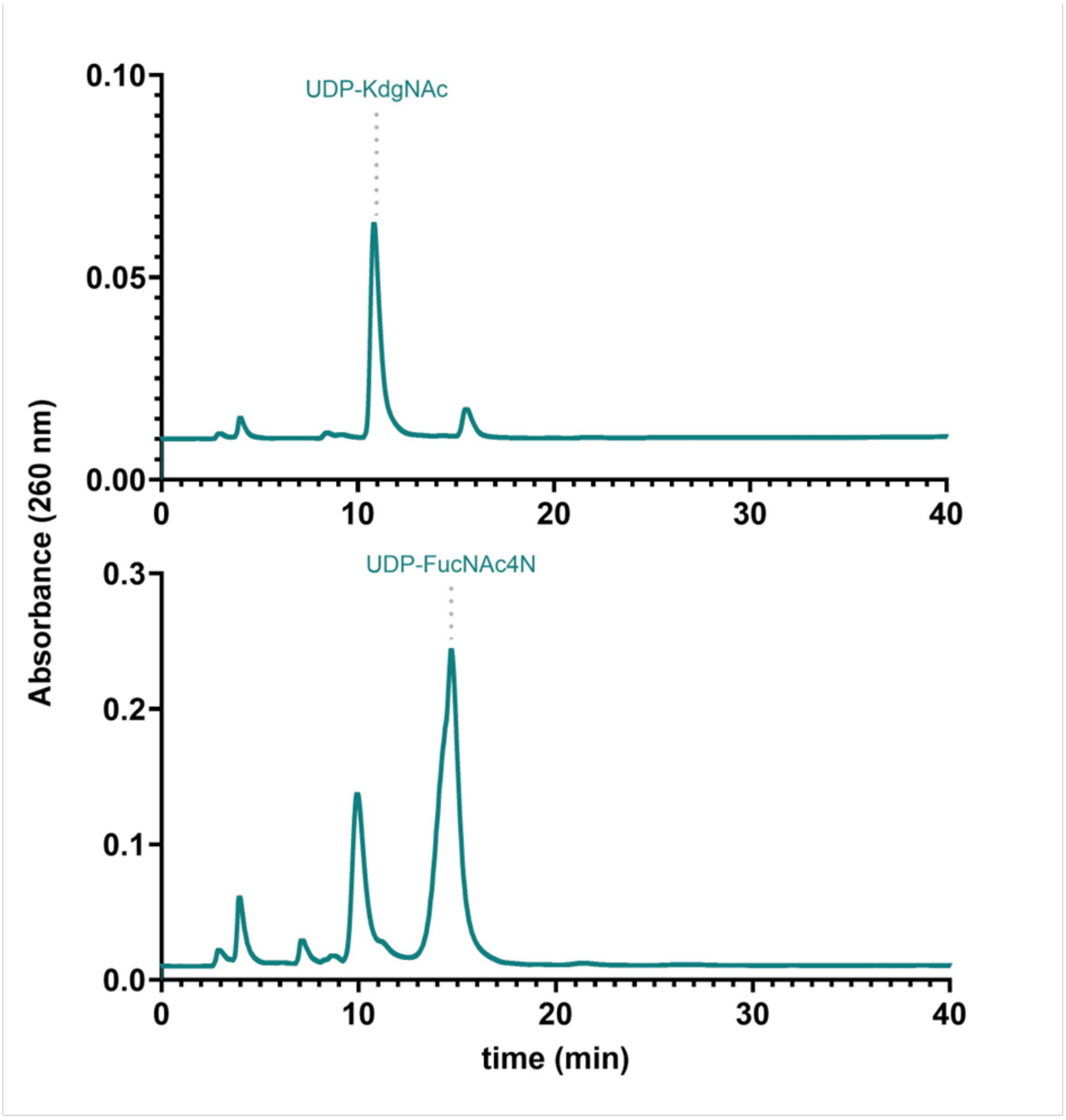


**B.**

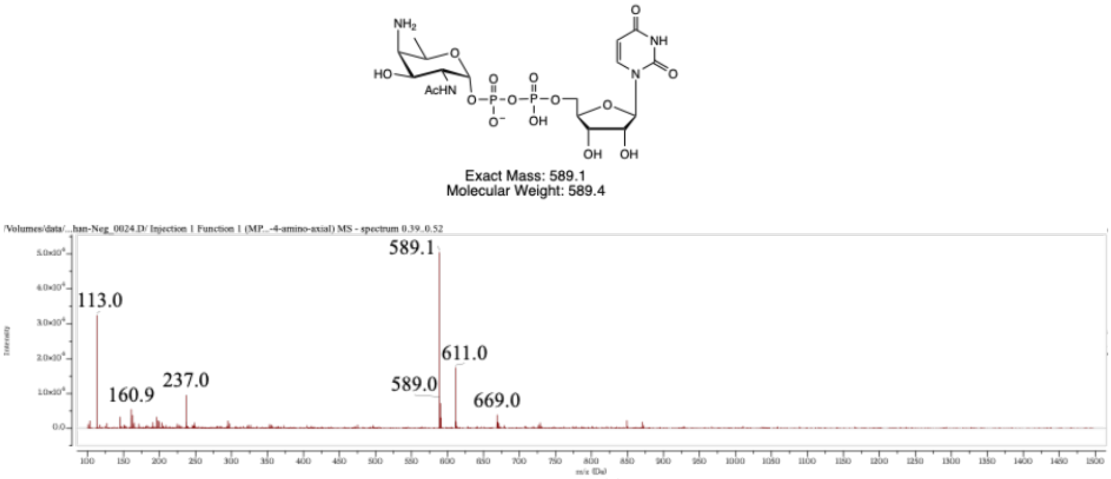


**Figure S6**: Characterization of UDP-FucNAc4N from chemoenzymatic synthesis. **A**. top: A 10 µL injection of WcfR reaction starting material by anion-exchange HPLC on gradient C, bottom: A 100 µL injection of WcfR reaction after incubation with gradient C. **B**. LRMS ESI(-) of WcfR reaction elution fractions and exact molecular weight predictions of the UDP-FucNAc4N, 611.0 m/z peak is thought to be the FucNAc4N sodium adduct.

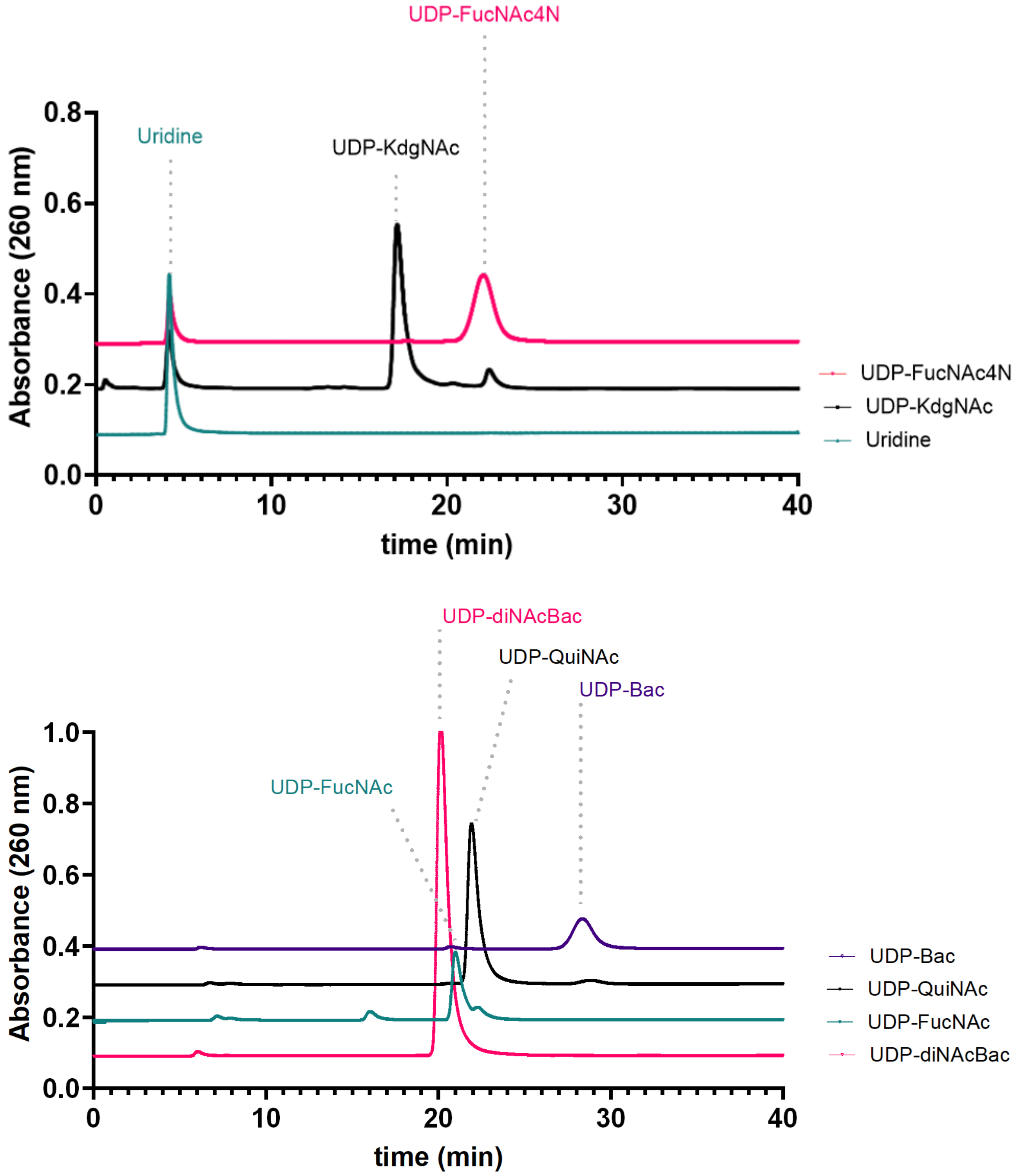


**Figure S7**: Calf Intestinal Alkaline Phosphatase (CIAP) treatment for removal of nucleotide contaminants. **Top**. A 200 µL injection of 500 µM CIAP treated UDP-FucNAc4N and UDP-KdgNAc and 5 µL injection of 9 mM uridine on Gradient C. **Bottom**. Injections of CIAP-treated materials, 250 µL of 1mM diNAcBac, 200 µL of 250 µM FucNAc, 250 µL of 500 µM of UDP-QuiNAc and 200 µL of 500 µM UDP-Bac on Gradient C.

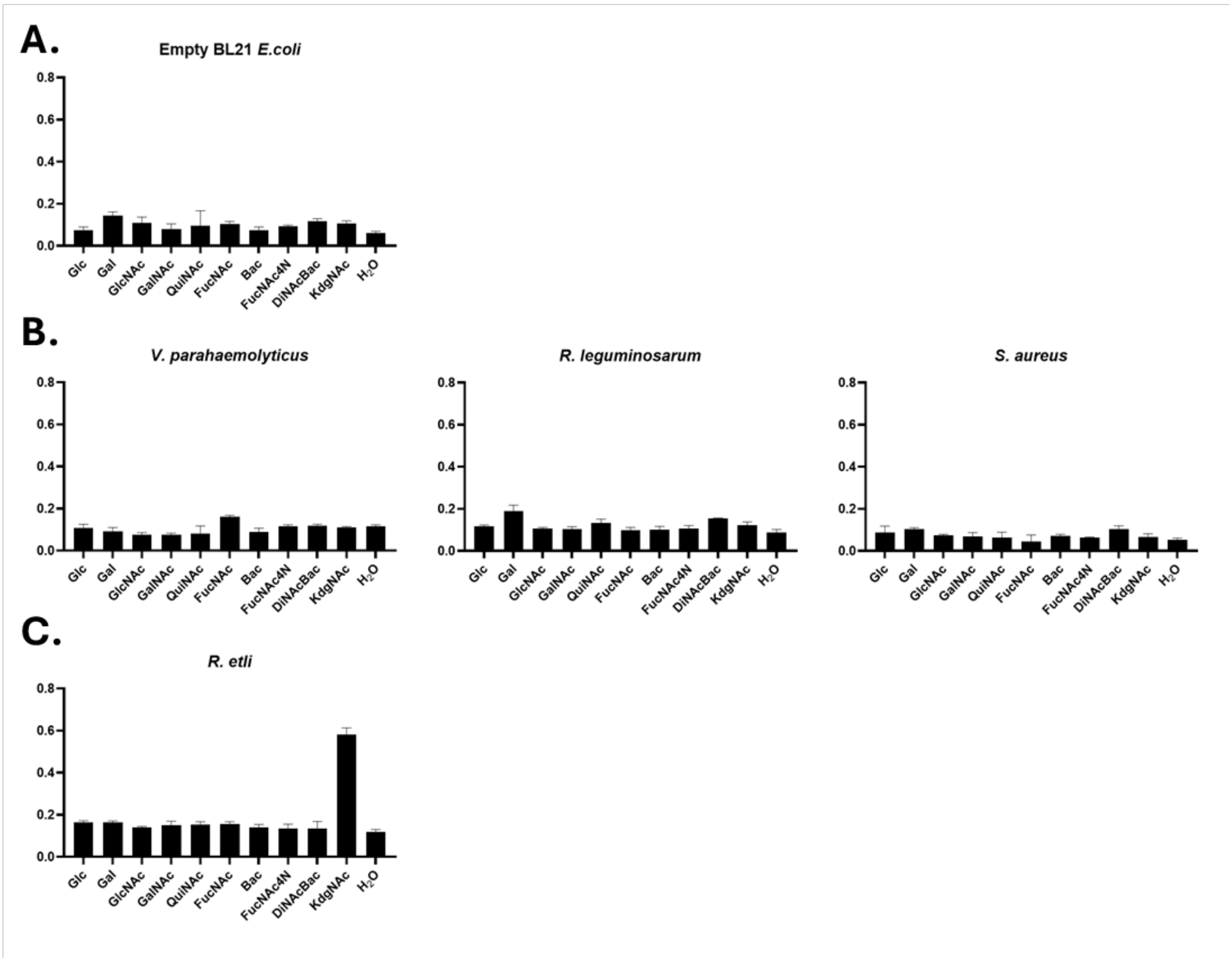


**Figure S8**: CEF-Glo control data. **A**. BL21 *E. coli* empty plasmid CEF controls. **B**. CEF-Glo screens of SmPGTs with no identified UDP-sugar substrate. **C**. Positive control with *R. etli* WreU.

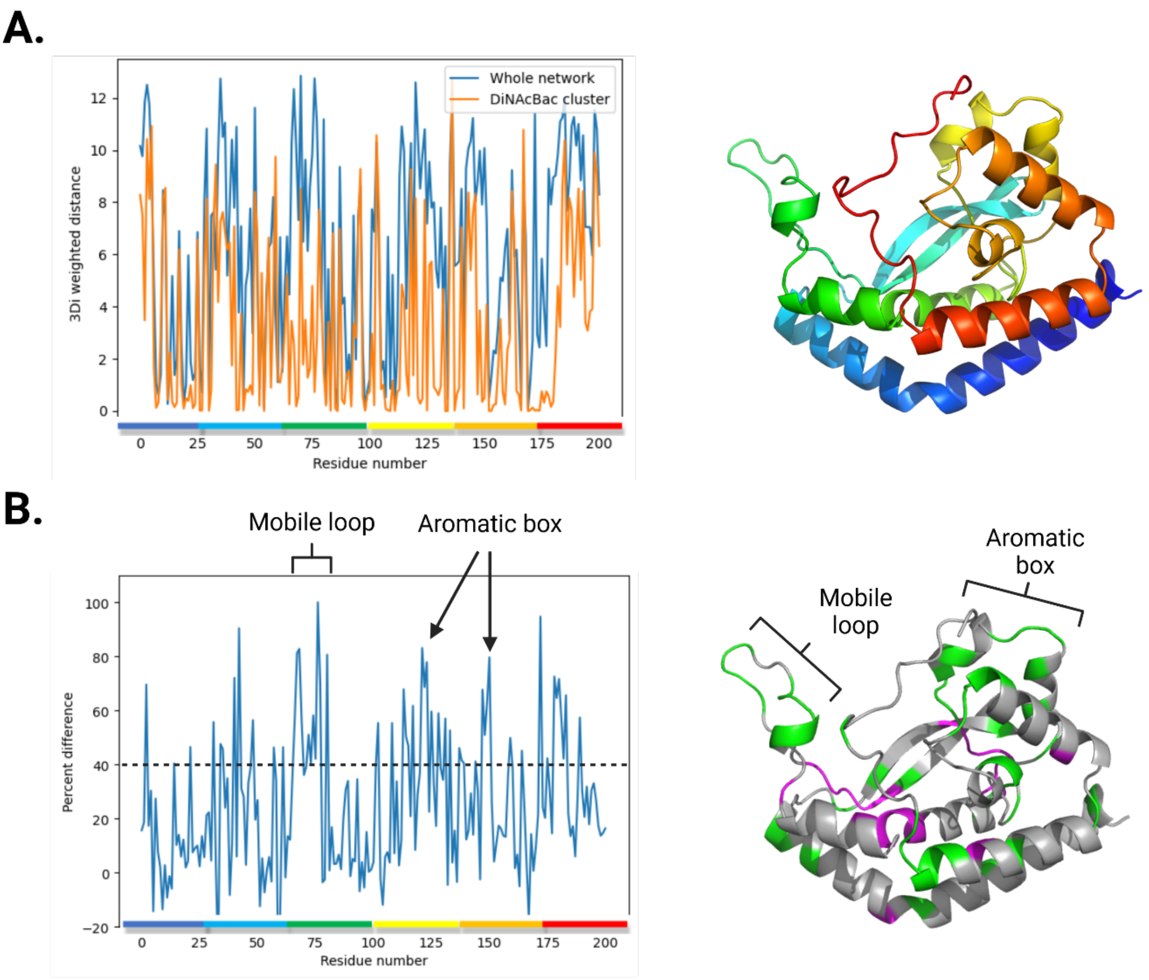


**Figure S9:** Example of structural conservation analysis for chosen smPGT. **A**. Euclidean dissimilarity for 3Di alignments at *C. concisus* PglC residues. Whole network alignment (blue) shows higher dissimilarity than the substrate cluster alignment (orange), particularly at prominent structural features like the mobile loop. Colors above residue numbers map to the protein structure on the right for visualization. **B**. Percent difference between curves in panel **A** highlights distinct substrate-specific structural features. Black dotted line shows 40% threshold used to color green residues on the lower right structure. Highlighted mobile loop and aromatic box correlate with previous studies on PglC (57, 58).

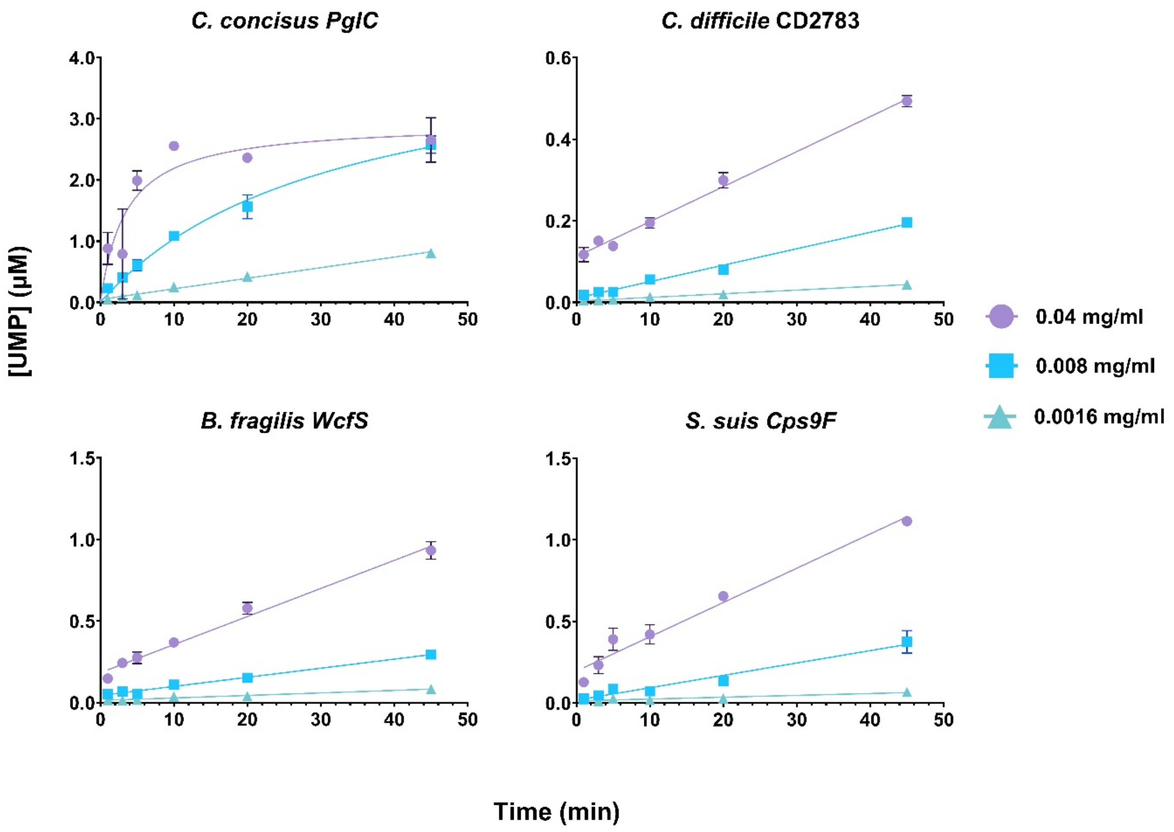


**Figure S10**: Kinetic analysis of selected SmPGTs in CEF.

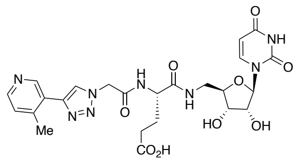

***Compound 1***

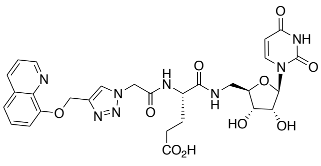

***Compound 2***

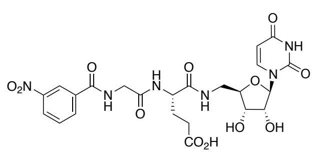

***Compound 3***

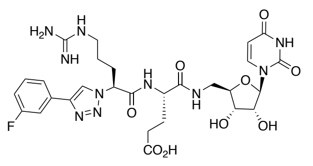

***Compound 4***

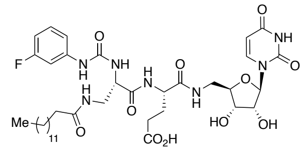

***Compound 5***

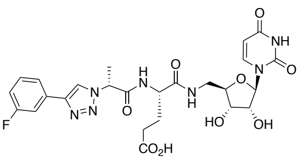

***Compound 6***

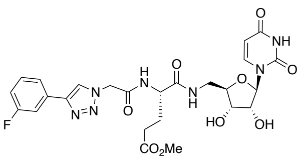

***Compound 7***

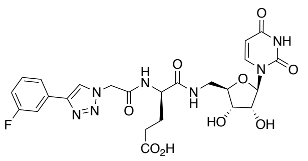

***Compound 8***

**Figure S11:** Structures of nucleoside analogs used for inhibitor screening.

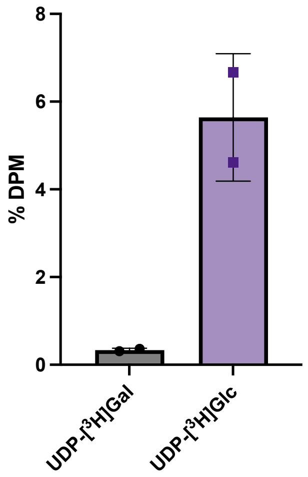


**Figure S12:** Radioactivity-based substrate assay with CEF containing overexpressed *R. leguminosarum* PssA. Assays were carried out as previously reported (59) with commercial UDP-[$^3$H]Hex sugar donors*.* Activity is reported as the percentage of disintegrations per minute (% DPM) in the organic layer normalized to the total disintegrations per minute per quench point. Error bars are given for mean ± standard deviation (SD), n = 2.

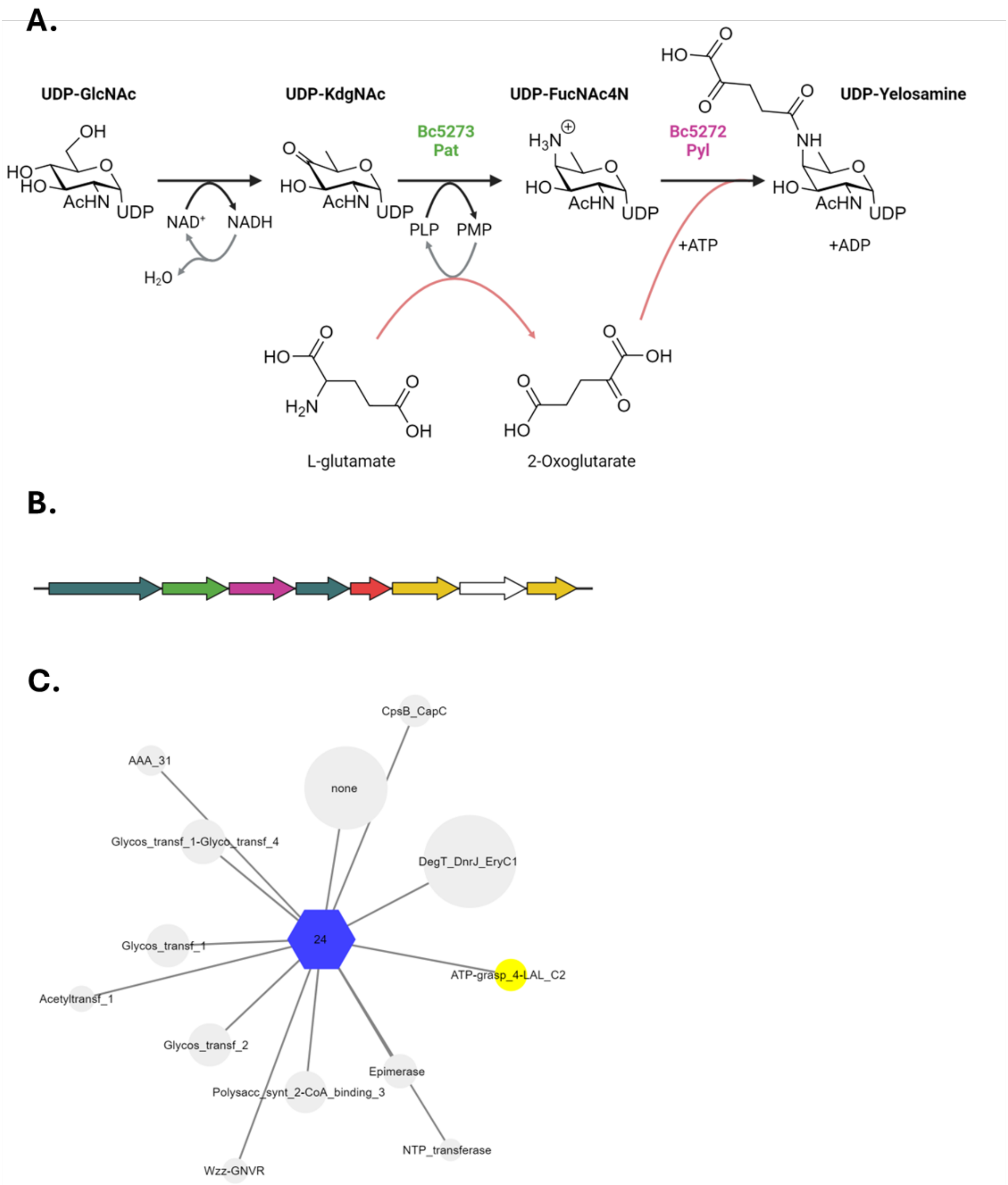


**Figure S13**: UDP-Yelosamine biosynthesis and operon analysis: **A**. Proposed biosynthetic route for generation of UDP-Yelosamine (47). **B**. Biosynthetic operon for secondary cell wall polysaccharide generation in *Bacillus cereus* ATCC 14579. SmPGT is shown in red, glycosyl transferases in yellow, UDP-biosynthesis enzymes in teal, with Pat and Pyl enzymes shown with same coloring as A., genes with unknown function are in white. **C**. Genome Neighbourhood Network of cluster 24, containing SmPGT shown in B., as hub-nodes and neighbouring Pfam familys as spoke-nodes. Highlighted in yellow is the frequent occurrence of the ATP-grasp (ATP-grasp_4-LAL_C2) domain protein (Pyl) within the cluster. The DegT_DnrJ_EryC1 aminotransferase Pfam family, of which Pat is a member, is also highly represented.

**Supplementary Methods**

**Typical chemical synthesis scheme of nucleoside analogs**

**Uridine loading with BAL resin.**

5'-Aminouridine was attached to BAL resin using a modified protocol (60, 61). BAL resin (Advanced Chemtech, 100-200 mesh, 0.6–1.2 mmol/g, 1% DVB) was weighed into a reaction vessel containing a frit (Torviq 10 mL Luer Lock Fritted Syringe from Fisher, Catalog No. NC9299151). The resin was swelled in 1% AcOH/DMF (1 g resin/10 mL) for 1 h. The solvent was removed, and the resin was treated with a 0.2 M mixture of 2',3' acetal-protected 5'-amino uridine (62) (2 equiv) in 1% AcOH/DMF. The reaction was agitated for 1 h (ambient temperature). Then a mixture of $NaBH_3CN$ (2 equiv) in MeOH (0.45 M) was added directly to the slurry of resin and agitated overnight. *Note: A closed reaction vessel is not recommended as production of gas is observed after the addition of $NaBH_3CN$. After 18 h, the solvent mixture was evacuated and the resin was rinsed with DMF, $CH_2Cl_2$, MeOH, and $H_2O$ (2 x 5 mL each). The resin, in a slurry of $H_2O$, was frozen and lyophilized to provide the dried resin.

***3-NO₂Ph (Compound 3).*** *(S)-5-((((2R,3S,4R,5R)-5-(2,4-dioxo-3,4-dihydropyrimidin-1(2H)-yl)-3,4-dihydroxytetrahydrofuran-2-yl)methyl)amino)-4-(2-(3-nitrobenzamido)acetamido)-5-oxopentanoic acid.* Resin (150 mg, loading: 0.53 mmol/g, 0.08 mmol, 1 equiv) was weighed into a 10 mL fritted syringe. The resin was swelled for 15 min in $CH_2Cl_2$ (5 mL). Then a mixture of Fmoc-Glu(OtBu)-OH (5 equiv), HBTU (5 equiv), and Hünig's base (10 equiv) in DMF (5 mL) was added to the resin. The mixture was agitated for 2 h (ambient temperature). After this time, the solvent was removed, and the resin was rinsed with DMF (2 x). Then a mixture of 20% piperidine/DMF (v/v) was added to the resin and agitated for 20 min. The solvent was removed, and the resin was rinsed with DMF (2 x). A mixture of Fmoc-Gly-OH (5 equiv), HBTU (5 equiv), and Hünig's base (10 equiv) in DMF (5 mL) was added to the resin. The mixture was agitated for 2 h (ambient temperature). Then a mixture of 20% piperidine/DMF (v/v) was added to the resin and agitated for 20 min. The solvent was removed, and the resin was rinsed with DMF (2 x). Then a mixture of carboxylic acid (5 equiv), HBTU (5 equiv), and Hünig's base (10 equiv) in DMF (10 mL) was added to the resin. The mixture was agitated for 2 h (ambient temperature). After this time, the solvent was removed, and the resin was rinsed with DMF and $CH_2Cl_2$ (2 x each). The crude product was cleaved from the resin with 2 mL TFA/TIPS/$H_2O$ (95:2.5:2.5) for 2 h. The TFA cleavage was concentrated in volume under a stream of $N_2$ and the crude product was precipitated with –20 ºC diethyl ether. The resulting slurry was centrifuged, and the TFA/ether supernatant was decanted to yield the crude pellet. The pellet was resuspended in MeCN/$H_2O$ and purified by preparative RP-HPLC (Luna 5 µm $C_{18}$(2) 100 Å, 250 x 21.2 mm Phenomenex column) with a gradient of 20-75% B over 25 min, flow rate: 10 mL/min [solvents A: $H_2O$ (0.1% TFA), B: MeCN (0.1% TFA)]. The purified product was transferred to 50 mL centrifugation tubes, frozen in $LN_2$, and lyophilized to yield a fluffy white solid (18% yield, 8.2 mg).

**$^1$H NMR (600 MHz, DMSO-*d*$_6$)** δ 12.09 (s, 1H), 11.29 (d, *J* = 2.4 Hz, 1H), 9.20 (t, *J* = 5.8 Hz, 1H), 8.71 (t, *J* = 2.0 Hz, 1H), 8.40 (ddd, *J* = 8.2, 2.4, 1.1 Hz, 1H), 8.31 (dt, *J* = 7.7, 1.3 Hz, 1H), 8.22 (d, *J* = 8.2 Hz, 1H), 8.11 (t, *J* = 6.0 Hz, 1H), 7.78 (t, *J* = 8.0 Hz, 1H), 7.63 (d, *J* = 8.1 Hz, 1H), 5.72 (s, 1H), 5.61 (dd, *J* = 8.0, 2.2 Hz, 1H), 5.37 (d, *J* = 5.8 Hz, 1H), 5.15 (d, *J* = 5.1 Hz, 1H), 4.31 (td, *J* = 8.6, 5.1 Hz, 1H), 3.98 (dd, *J* = 24.6, 5.7 Hz, 3H), 3.88 – 3.80 (m, 2H), 3.48 – 3.37 (m, 1H), 2.23 (pt, *J* = 9.5, 5.2 Hz, 2H), 1.98 – 1.90 (m, 1H), 1.74 (dtd, *J* = 14.6, 9.3, 5.7 Hz, 1H).

**$^{13}$C NMR (151 MHz, DMSO)** δ 173.9, 171.4, 168.8, 164.6, 163.0, 150.7, 147.8, 141.1, 135.4, 133.8, 130.1, 126.0, 122.1, 102.0, 87.9, 82.6, 72.5, 70.6, 51.9, 42.8, 40.9, 30.1, 27.4.

**HPLC R$_t$:** 10.8 min (260 nm).

**HRMS (ESI$^+$)** m/z: [M+H]$^+$ Calc'd for $C_{23}H_{27}N_6O_{12}^+$ 579.1681; found 579.1690.

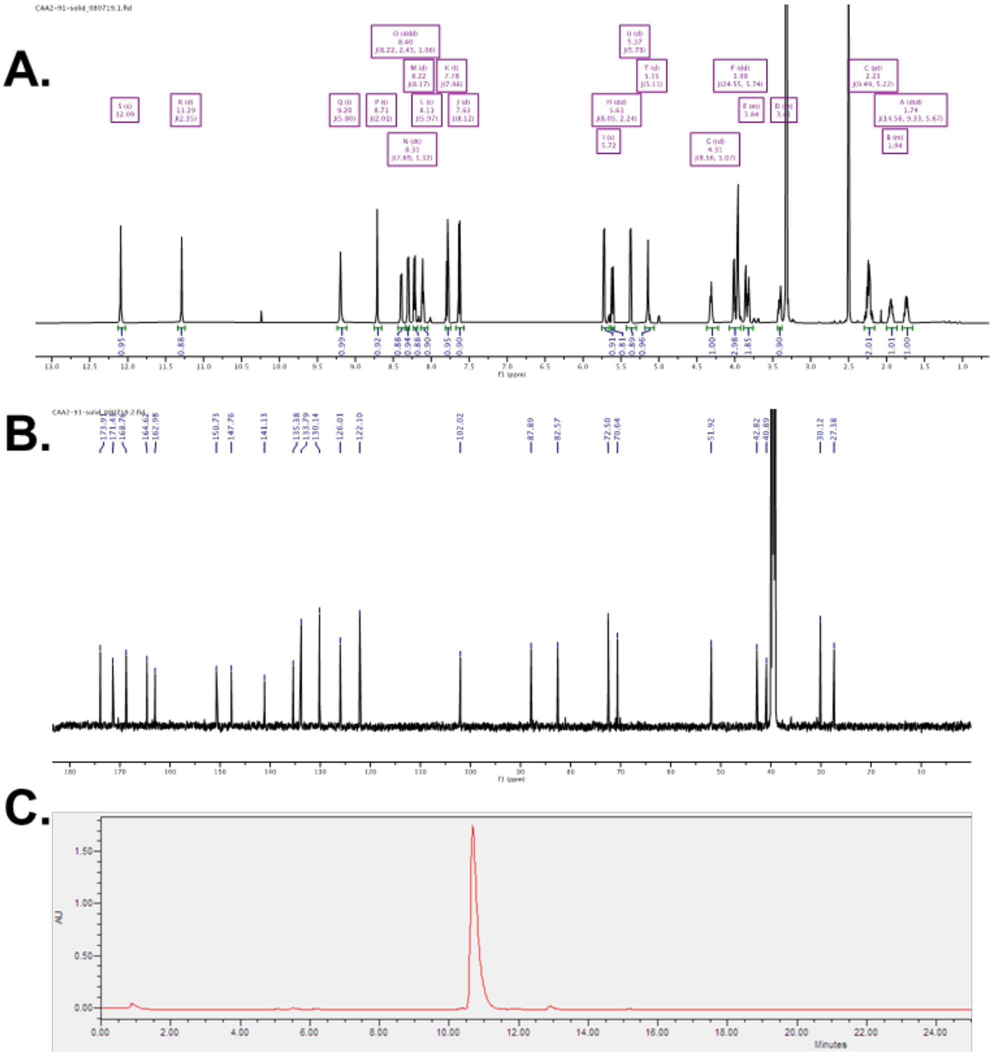

Characterization of compound **3**. **A:** $^1$H NMR spectrum (600 MHz, DMSO-d$_6$) of compound **3**. **B:** $^{13}$C NMR spectrum (151 MHz, DMSO- d$_6$) of compound **3**. **C:** Crude RP-HPLC UV (260 nm) spectrum of compound **3**.
